# Malate-dependent Fe accumulation is a critical checkpoint in the root developmental response to low phosphate

**DOI:** 10.1101/095497

**Authors:** Javier Mora-Macías, Jonathan Odilón Ojeda-Rivera, Dolores Gutiérrez-Alanís, Lenin Yong-Villalobos, Araceli Oropeza-Aburto, Javier Raya González, Gabriel Jiménez-Domínguez, Gabriela Chávez-Calvillo, Rubén Rellán-Álvarez, Luis Herrera-Estrella

**Affiliations:** Laboratorio Nacional de Genómica para la Biodiversidad (Langebio)/Unidad de Genómica Avanzada, Centro de Investigación y Estudios Avanzados del Instituto Politécnico Nacional, 36500 Irapuato, Guanajuato, México

## Abstract

Low phosphate (Pi) availability constrains plant development and crop production in both natural and agricultural ecosystems. When Pi is scarce, modifications of root system architecture (RSA) enhance soil exploration ability and can lead to an increase in Pi uptake. In *Arabidopsis*, an iron-dependent determinate developmental program that induces premature differentiation in the root apical meristem (RAM) begins when the root tip contacts low Pi media, resulting in a short-root phenotype. However, the mechanisms that enable the regulation of root growth in response to Pi-limiting conditions remain largely unknown. Cellular, genomic and transcriptomic analysis of low-Pi insensitive mutants revealed that the malate-exudation related genes *SENSITIVE TO PROTON RHIZOTOXICITY* (*STOP1*) and *ALUMINUM ACTIVATED MALATE TRANSPORTER 1* (*ALMT1*) represent a critical checkpoint in the root developmental response to Pi starvation in *Arabidopsis thaliana*.

## Introduction

Phosphorus is an essential nutrient for plant development, a constituent of key molecules such as nucleic acids, ATP and membrane phospholipids. Plants take up and metabolize phosphorus in the inorganic form of orthophosphate (Pi) (1). Pi is the least accessible macronutrient in many natural and agricultural ecosystems and its low availability in the soil often limits plant growth and productivity. Under phosphate limiting conditions (-Pi), plants activate an array of genetic (2–6), biochemical (7, 8) and morphological modifications (9–11) that enhance their ability to cope with low Pi availability.

*Arabidopsis* responses to low Pi availability have been divided in systemic responses that depend upon the internal Pi status of the plant and local responses that depend upon the level of Pi available in the external medium (5, 12). A molecular dissection of local and systemic responses to Pi starvation using transcriptomic analysis has been reported (5). Systemic responses include the upregulation of genes involved in the overall enhancement of Pi uptake and Pi internal use efficiency, and are largely controlled by the master regulator PHR1 (a Myb transcription factor) (6, 13). Local responses include alterations of root traits such as the inhibition of primary root growth (14), an increase in lateral root density (10) and higher density and length of root hairs (9). These changes in root system architecture have been proposed to enhance the soil exploration ability of the plant by increasing root surface area of exploration in the top layers of the soil where Pi tends to accumulate. Some evidence indicates that there is some degree of crosstalk between local and systemic responses to low Pi availability as mutants altered in root system architecture responses to low Pi also have an altered expression of genes involved in systemic responses (15).

Growth of *Arabidopsis* seedlings under *in vitro* Pi-limiting conditions induces a determinate developmental program known as RAM exhaustion (11). RAM exhaustion consists of the loss of meristematic potential and the arrest of cell proliferation, leading to the inhibition of primary root growth. Meristematic potential is lost due to premature differentiation of the cells that constitute the stem cell niche (SCN), which includes the quiescent center (QC). Cell proliferation in the RAM is lost due to a gradual reduction in mitotic activity. Root-tip contact with low phosphate media (16) in the presence of iron (17) has been reported as essential for the inhibition of primary root growth in response to Pi deficiency conditions in *Arabidopsis thaliana. Arabidopsis* mutants that fail to trigger root system morphological responses to low Pi have been reported previously (15, 16). Two of these mutants, *low phosphate root 1* and *2* (*lpr1 and lpr2*) were found to be affected in genes encoding multicopper oxidases, suggesting that a metal with different levels of oxidation could be involved in the alteration of root system architecture in response to low Pi availability (16). A mutant that is hypersensitive to low Pi, *phosphate deficiency response 2* (*pdr2*), and triggers the root system response to low Pi availability faster that the wild-type (WT) was also reported (18). *PDR2* encodes an ER localized P_5_-type ATPase(19). *PDR2* and *LPR1* have been proposed to orchestrate RAM-exhaustion in a genetically interacting route under low Pi conditions (19). Whilst the precise function of *PDR2* has not been determined, LPR1 is essential for primary root inhibition under low Pi conditions and it has been shown to have ferroxidase activity (16, 19, 20). An LPR1-dependent accumulation of Fe^3+^ in the apoplast of cells in the elongation and meristematic regions of the primary root, that triggers the production of reactive oxygen species (ROS), is essential for meristem exhaustion in low-Pi media (20). ROS generation correlates with callose deposition in the RAM, which was proposed to activate meristem exhaustion by blocking the cell-to-cell movement of SHORT-ROOT (SHR), a transcription factor that is essential for stem cell maintenance in the RAM (20). However, the mechanism that regulates iron accumulation and relocation in the RAM remains largely unknown.

In this work, we characterized two <u>***L***</u>*ow* ***<u>p</u>****hosphate* ***<u>i</u>****nsensitive* mutants (*lpi5 and lpi6*), which, in contrast to the short root phenotype of WT *Arabidopsis* seedlings, show normal primary root elongation in low Pi media. Mapping by sequencing revealed that the *lpi5* corresponds to *SENSITIVE TO PROTON RHIZOTOXICITY* (*STOP1*) and *lpi6* to *ALUMINUM ACTIVATED MALATE TRANSPORTER 1* (*ALMT1*), the two genes previously reported to be responsible for activating malate efflux in the roots of Arabidopsis seedlings exposed to toxic concentrations of aluminum (21-23). Genetic, cellular and transcriptomic analysis shows that *STOP1* and *ALMT1* are required for a malate-dependent accumulation of iron in the root meristem, which leads to alterations in the redox balance that triggers primary root growth inhibition and RAM exhaustion in response to Pi deficiency conditions in *Arabidopsis thaliana*.

## Results

### *Arabidopsis* EMS-mutants *lpi5* and *lpi6* show indeterminate primary root growth under phosphate deficiency conditions

The *Arabidopsis thaliana* Col-0 accession seedlings grown in media containing low Pi concentrations (below 50 μM Pi) show a short root phenotype defined by a determinate pattern of primary root growth and RAM differentiation. To identify mutants that are insensitive to the effect of low Pi on primary root growth, we screened an EMS-mutagenized Col-0 population for mutant lines presenting a long root phenotype under low-Pi (-Pi, 5 μM Pi) conditions. We isolated approximately 50 mutant lines with different alterations in the primary root growth in response to low-Pi availability. Two lines that were insensitive to the effect of low Pi on primary root growth, which we named *low phosphorus insensitive 5 and low phosphorus insensitive 6* (*lpi5* and *lpi6*), were chosen for further characterization (Figure 1A, B, C). When grown under high Pi (+Pi; 100μM Pi) conditions, both *lpi5* and *lpi6* seedlings presented a primary root length similar to the WT *Arabidopsis* Col-0 accession (Figure 1A). Under -Pi conditions at 10 dag, WT plants had a visible reduction in primary root growth, whereas *lpi5* and *lpi6* seedlings had primary root elongation similar to that of WT seedlings grown in +Pi media which is four-fold longer than that of Pi-deprived WT seedlings (Figure 1B). Although *lpi5* seedlings had a long root phenotype in -Pi media, they had a slightly shorter primary root than the WT and *lpi6* in +Pi media and were also slightly shorter than *lpi6* in -Pi medium (Figure 1A, C). Analysis of segregation frequencies under -Pi conditions showed that the long root phenotype in *lpi5 and lpi6* is the result of single recessive mutations (Supplementary Table 1).

In Pi-deprived *Arabidopsis* seedlings, the short-root phenotype is accompanied by a reduction in cell proliferation and meristematic activity during the process of RAM exhaustion (11). To test for signs of RAM exhaustion in *lpi5* and *lpi6* seedlings grown in -Pi media, we examined the expression of the *proCycB1::GUS* cell cycle activity marker (24) and the *proQC46::GUS* quiescent center (QC) identity marker (25). In +Pi media, WT, *lpi5* and *lpi6* seedlings showed clear cell proliferation activity as indicated by the *proCycB1::GUS* signal and an active QC as shown by the *proQC46::GUS* signal (Figure 1D). In -Pi media at 10 dag the cell cycle and QC marker genes were undetectable in WT seedlings, whereas GUS staining was clearly detectable for both the cell cycle (*proCycB1::GUS*) and QC (*proQC46::GUS*) markers in the primary root of *lpi5* and *lpi6* (Figure 1D). These results show that, in contrast to the WT, low Pi does not trigger meristem exhaustion in the RAM of *lpi5* and *lpi6* seedlings.

**Figure 1.**
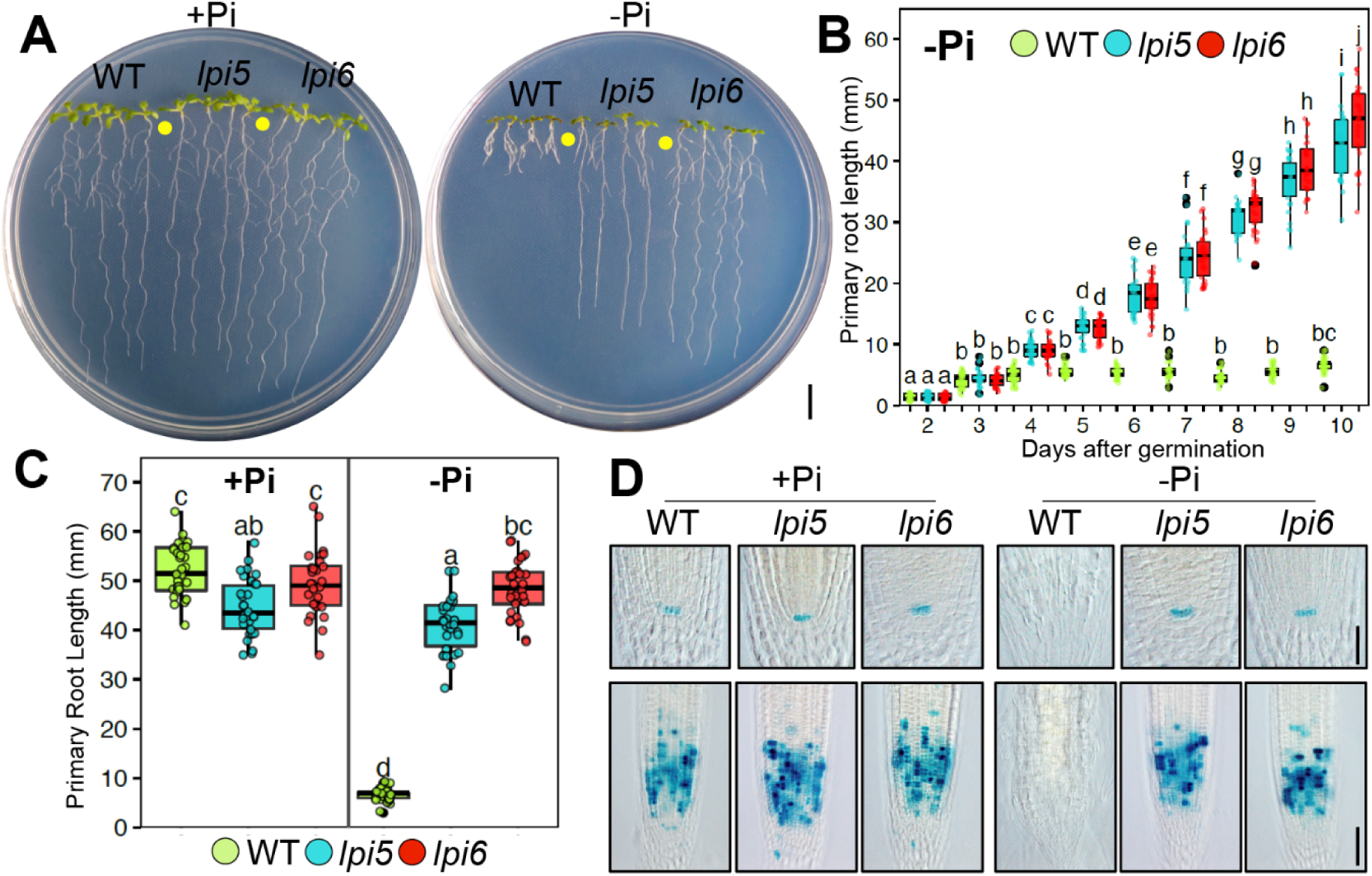
Low-Pi insensitive mutants continue primary root growth and RAM maintenance under Pi deficiency conditions. (A) Phenotypes of WT, lpi5 and lpi6 seedlings grown under high (100 μM Pi; +Pi) and low Pi (5 μM Pi; -Pi) conditions 10 days-after-germination (dag). Scale bar equals 1 centimeter (cm). (B) Primary root growth kinetics of seedlings grown under -Pi conditions from 2 to 10 dag. Green, blue and red dots depict WT, *lpi5* and *lpi6* individuals (n=30 from 3 independent experiments), respectively. Statistical groups were determined using a Tukey HSD test (P-value < .05) and are indicated with letters. (C) Primary root length of seedlings grown under +Pi and -Pi conditions. Green, blue and red dots depict WT, lpi5 and lpi6 individuals (n=30 from 3 independent experiments), respectively. Statistical groups were determined using a Tukey HSD test (P-value < .05) and are indicated with letters. (D) GUS staining of *proCycB1::GUS* and *proQC46::GUS* expression in the RAM of WT, *lpi5* and *lpi6* seedlings 10 dag. Scale bar indicates 100 μm.

### *lpi5* and *lpi6* have mutations in the transcription factor *STOP1* and the malate transporter *ALMT1*, respectively

We used a mapping-by-sequencing approach to identify the genes responsible for the *lpi5* and *lpi6* phenotypes (See Materials and Methods). We identified 12 and 18 specific homozygous variants (missense, frameshift and splice-donor) potentially linked to *lpi5* and *lpi6* mutant phenotypes, respectively. Given the alterations of primary root of *lpi5* and *lpi6* seedlings in response to an environmental stress, we focused on the homozygous mutations that could potentially be linked to alterations in root morphological or root responses to abiotic stress. Among the potential candidates responsible for the root phenotype of *lpi5* and *lpi6* under low Pi conditions, *SENSITIVE TO PROTON RHIZOTOXICITY* (*STOP1*) and *ALUMINUM ACTIVATED MALATE TRANSPORTER 1* (*ALMT1*) were particularly interesting because both genes have been previously reported to participate in the tolerance of the *Arabidopsis* root to toxic concentrations of Al^+3^. *STOP1* (At1g34370) encodes a zinc finger protein transcription factor that plays a critical role in Al^3+^ tolerance and acid soil tolerance in *Arabidopsis* (21). *ALMT1* encodes a transmembrane protein that mediates malate efflux in the root in response to the presence of toxic Al^3+^ levels (22), and its expression has been shown to be activated by *STOP1* in response to Al stress conditions (21).

To test whether the long root phenotype of *lpi5, lpi6* was indeed due to lesions in STOP1 and ALMT1, we tested the phenotype of T-DNA insertional mutants in *STOP1 (SALK_114108)* and *ALMT1 (SALK_009629)* in -Pi and +Pi media. In +Pi media, WT, *lpi6* and *almt1* seedlings had a similar primary root length, whereas *lpi5* and *stop1* had a slightly shorter root length than the WT (Figure 2A-B). In -Pi media, the T-DNA insertional mutants *stop1* and *almt1* had a long root phenotype that contrasted with the short root phenotype of the WT (Figure 2A-B). It has been reported that *stop1* is sensitive to low pH (4.7) and levels of Al^3+^(2 μM) that are not yet toxic for *Arabidopsis* WT seedlings and that *almt1* is also sensitive to this Al^3+^ concentration (28). We also found that *lpi5* was sensitive to low pH and Al (as is *stop1)* and that *lpi6* was sensitive to Al but not to low pH (as observed for *almt1)* (Supplementary Figure 1). Crosses between *lpi5* and *almt1*, and between *lpi6* and *stop1*, showed that *lpi5* and *lpi6* are mutant alleles of *STOP1* and *ALMT1*, respectively (Supplementary Figure 1).

Mapping-by-sequencing of *lpi5* revealed a C:T (CAT:TAT) substitution in the +84 position of *STOP1. In silico* analysis predicted that the *lpi5* mutation caused an amino acid substitution (H168Y) that replaces one of the two histidine residues that are crucial for the formation of first of the four zinc fingers of the DNA binding domain of STOP1 (Figure 2C). The first zinc finger domain of STOP1 is critical to bind to the promoter of its target genes (29). Mapping-by-sequencing of the *lpi6* mutant revealed a 13-base-pair deletion in the first exon of *ALMT1* starting at position +757 (Figure 2D). *In silico* analysis predicted that this deletion causes a frameshift mutation that produces an aberrant protein that lacks 200 amino-acids of the carboxy-terminal moiety of ALMT1 (Figure 2D). Our *in silico* sequence analysis further confirmed that *lpi5* and *lpi6* are EMS-induced mutant alleles of *STOP1* and *ALMT1*, respectively.

**Figure 2.**
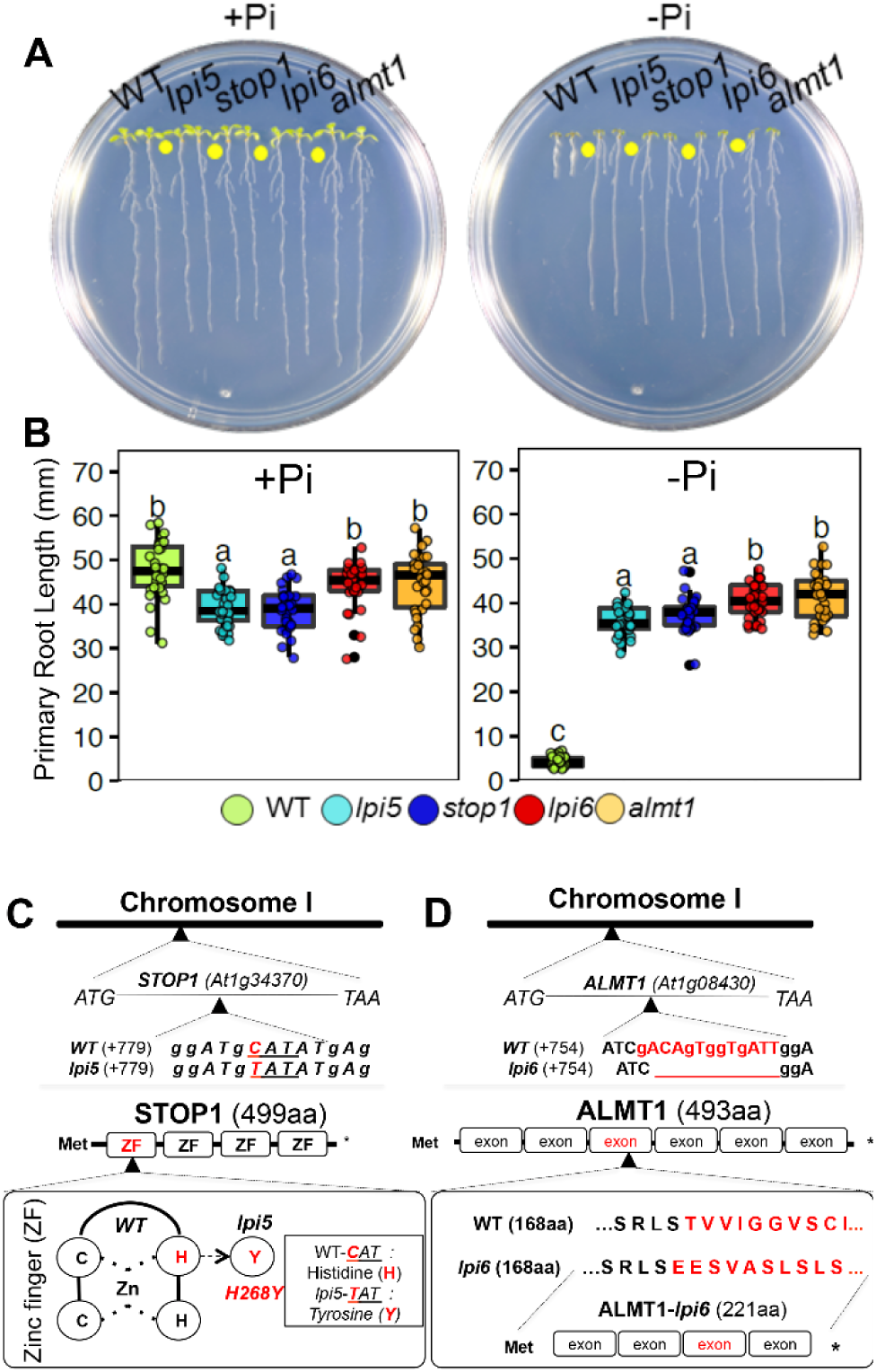
Mapping by sequencing revealed *lpi5 and lpi6* to be *stop1* and *almt1* mutants, respectively. (A) Phenotypes of Col-0, *lpi5, stop1, almt1* and *lpi6* seedlings grown under high phosphate (+Pi) and low phosphate (-Pi) conditions 10 dag. (B) Primary root length of WT, *lpi5, stop1, almt1* and *lpi6* seedlings grown under high phosphate (+Pi) or low phosphate (-Pi) conditions 10 dag. Dots depict WT, *lpi5, stop1, almt1* and *lpi6* individuals (n=30 from 3 independent experiments). Statistical groups were determined using a Tukey HSD test (P-value < .05) and are indicated with letters. (A) C:T (CAT:TAT) substitution in the +84 base within the At1g34370 locus of the *lpi5* genome. *lpi5* mutation causes an amino acid substitution (H168Y) that replaces one of the two histidine residues of the four zinc fingers that constitute STOP1 (B) 13 base pair deletion in the +757 base pair position within the first exon of the At1g08430 locus of the *lpi6* genome. The 13 base deletion present in *lpi6* causes a frameshift mutation that produces an aberrant protein with 200 amino-acids less than the WT.

### *ALMT1* is expressed in the RAM of *Arabidopsis* under Pi deficiency conditions before meristematic exhaustion

To determine whether the expression of *STOP1* and *ALMT1* is regulated by Pi availability in the root tip, we analyzed the expression of *STOP1* and *ALMT1* in the root apex of 5 dag WT and *stop1* seedlings using qRT-PCR (Figure 3A, Supplementary Figure 3). In the WT, *ALMT1* expression increased by approximately 4-fold in response to -Pi conditions, while the level of expression of *STOP1* was not significantly altered by Pi availability (Figure 3A). We also found that in the *stop1* background the expression of *ALMT1* was undetectable (Supplementary Figure 2). Our results confirm a previous report (21) showing that *STOP1* is essential for the expression of *ALMT1*, but also show that *STOP1* is required for the induction of *ALMT1* in response to low Pi.

Since *STOP1* and *ALMT1* appear to be essential for RAM-exhaustion under Pi deficiency conditions, we examined the cell-specific expression pattern of these two genes in seedlings grown under +Pi and -Pi conditions 5 dag, a time point prior to full RAM exhaustion (Figure 1C), but when primary root growth inhibition has already started (Figure 1C). Confocal microscopy of *proSTOP1::GUS::GFP* seedlings grown in +Pi media revealed that *STOP1* is expressed in the QC, columella, lateral root cap and epidermis (Figure 3B) and that its expression pattern is not altered in -Pi media (Figure 3B). On the other hand no detectable reporter activity of *proALMT1::GUS::GFP* was found in the RAM of seedlings grown under +Pi conditions, whereas in -Pi media expression of the reporter gene was clearly detectable in the proximal region of the SCN (QC, cortex and stele initials), vascular bundle, pericycle, cortex, endodermis, columella and lateral root cap (Figure 3B).

**Figure 3.**
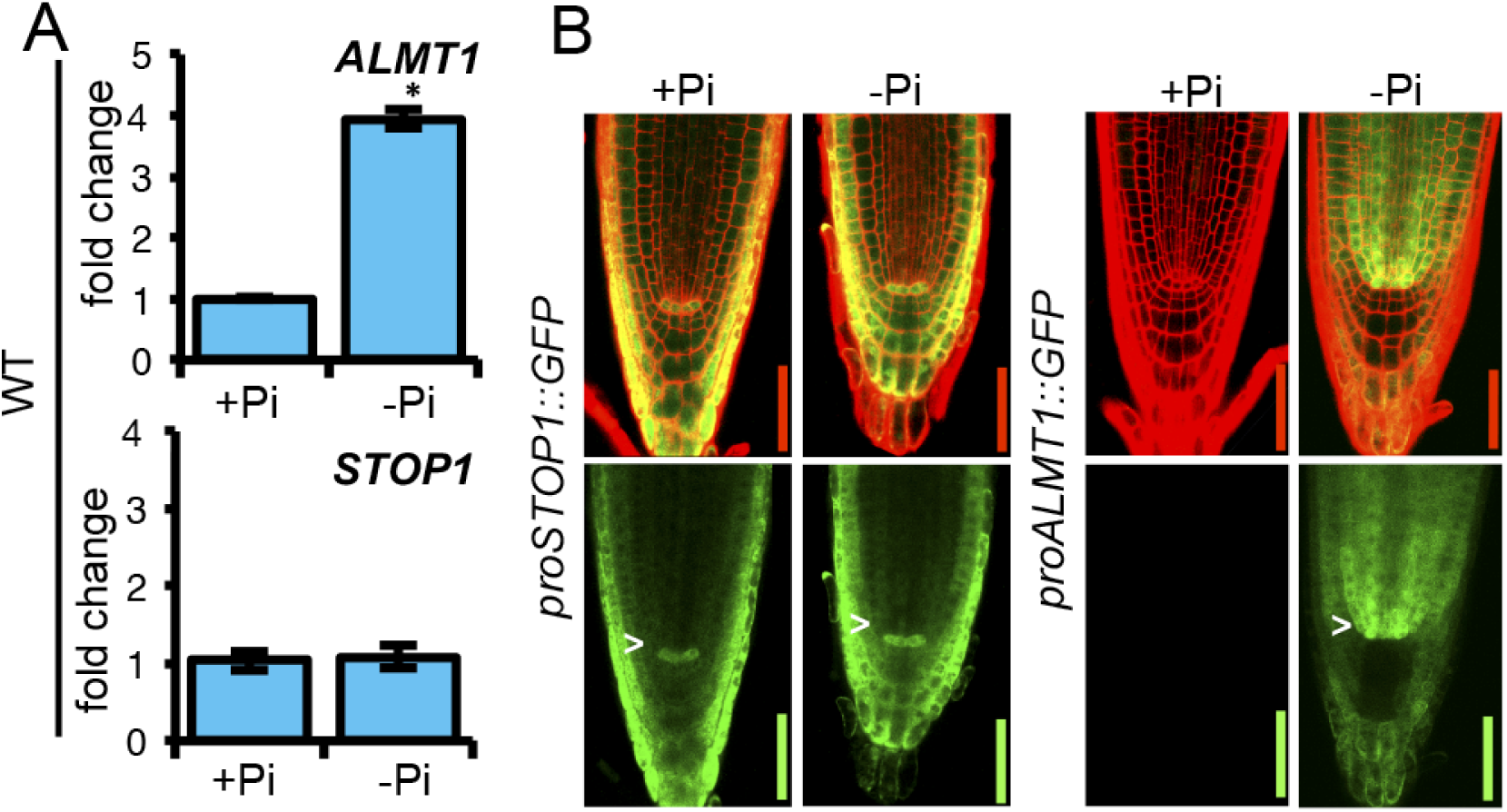
*STOP1* and *ALMT1* are expressed in the RAM of *Arabidopsis* under Pi deficiency conditions. (A) qRT-PCR analysis of *STOP1* and *ALMT1* expression in the root apex (2-3 mm) of WT (Col-0) plants. Bars represent the mean fold change ±SEM of 2 biological replicates with 3 technical replicates. WT +Pi samples were used as calibrator values. *ACT2* and *UBQ10* were used as internal controls. Asterisk indicates that the expression was significantly different between +Pi and -Pi conditions (Student t-test; p <0.05) (B) Transgenic Col-0 plants harboring transcriptional gene fusions containing the *STOP1* and *ALMT1* promoter fused to a double GFP-GUS reporter gene, respectively (*proSTOP1::GUS::GFP* and *proALMT1::GUS::GFP*), were grown under +Pi and -Pi conditions and expression activity was observed at 5 dag using confocal microscopy. Scale bar indicates 100 μm.

### Malate treatment rescues the determinate developmental program in the primary root of *stop1* and *almt1* in response to low Pi conditions

Since malate efflux has been shown to be affected in *stop1* and *almt1* mutants and both *stop1* and *almt1* present determinate primary root growth under Al^3+^ toxicity conditions (21, 22), we sought to determine whether malate exudation also played a role in the *Arabidopsis* primary root response to low Pi availability. To this end, we added increasing concentrations of malate to both +Pi and -Pi media and tested the effect of malate treatment on the primary root growth of WT, *stop1* and *almt1* seedlings (Figure 4 and Supplementary Figure 3). We observed that primary root growth was not altered by malate treatment under +Pi conditions in any of the tested lines (Supplementary Figure 3). However, in -Pi media, treatment with malate restored the short-root phenotype in *stop1* and *almt1* seedlings in a concentration-dependent manner (Figure 4A-B). Although *stop1* seedlings treated with 1 mM malate had significantly shorter roots than in media lacking malate, their primary roots were slightly, but statistically significantly, larger than those of the WT and *almt1* seedlings grown in the same media (Figure 4A-B). Malate treatment of Pi-deprived WT seedlings showed a small effect at 0.1 mM, however, this effect was not observed at higher malate concentrations (Figure 4A-B) as WT seedlings remained short under all treatments (Figure 4A).

To determine whether malate treatment activates RAM exhaustion in *stop1* and *almt1* seedlings, we examined the expression of *proCycB1::GUS* and *proQC46::GUS* reporter genes in Pi-deprived/malate-treated *stop1* and *almt1* seedlings. Clear signs of cell differentiation in the RAM of Pi-deprived/malate-treated *almt1* seedlings were observed and *proCycB1::GUS* and *proQC46::GUS* reporter activity was undetectable. In the case of Pi-deprived/malate-treated *stop1* seedlings, although cell proliferation was reduced, it was not completely arrested and *proQC46::GUS* expression was still clearly detectable (Figure 4C). No expression was found in the WT either in low-Pi media or low-Pi media supplemented with 1 mM malate (Figure 4C).

**Figure 4.**
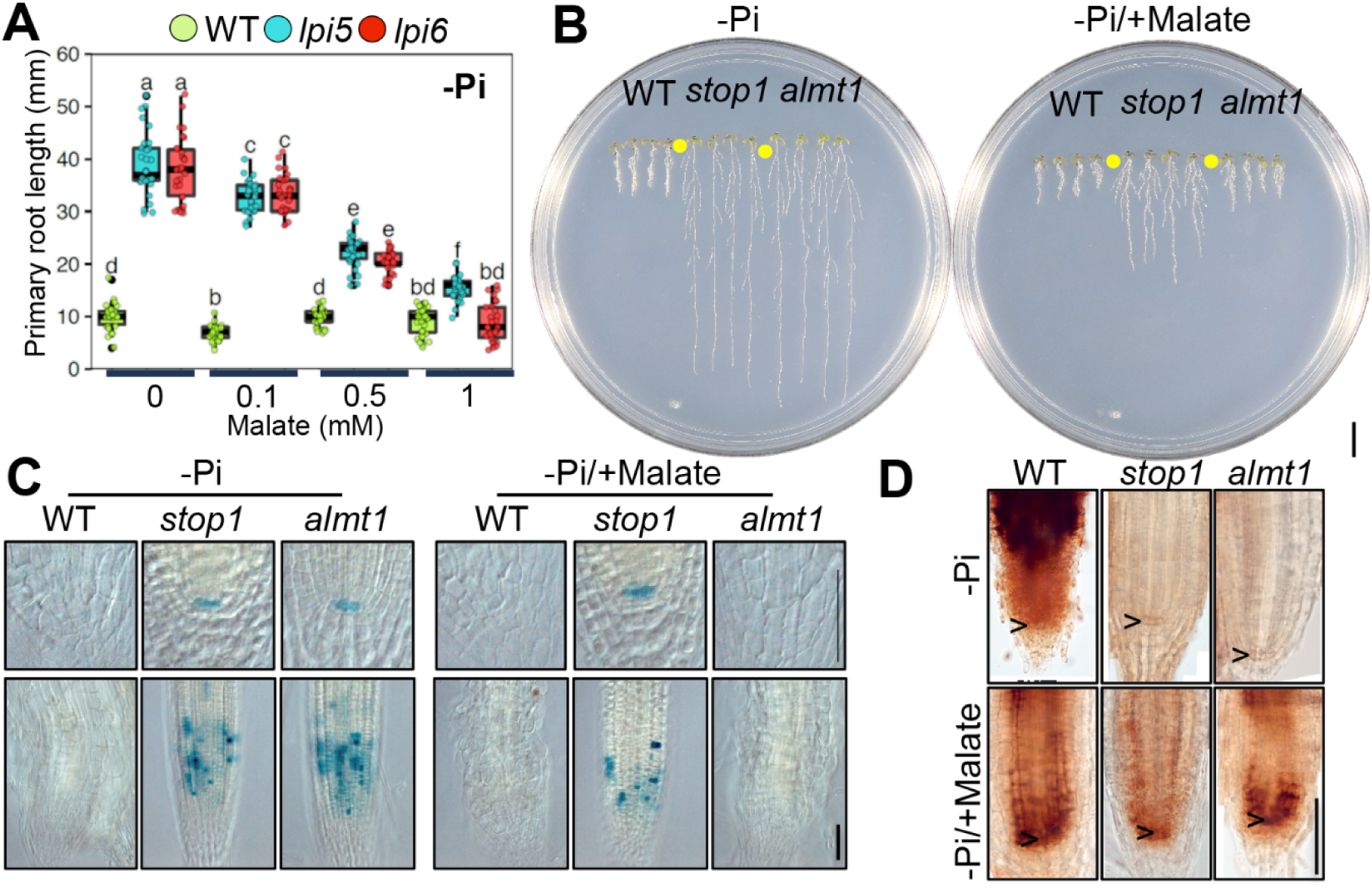
Malate treatment rescues the mutant phenotype of *stop1* and *almt1* seedlings. (A) Primary root length of 10 dag WT, *stop1* and *almt1* seedlings grown under increasing concentration of malate supplemented to -Pi medium. Green, blue and red dots depict WT, *stop1* and *almt1* individuals (n=30 from 3 independent experiments), respectively. Statistical groups were determined using Tukey HSD test (P-value <.05) and are indicated with letters. (B) Phenotypes of 10 dag WT, *stop1* and *almt1* seedlings grown under low phosphate medium (-Pi) and low phosphate medium supplemented with 1mM malate (-Pi/+Malate). Scale bar equals 1 mm. (C) *proCycB1::GUS* (lower panels) and *proQC46::GUS* (upper panels) expression (D) Perls-DAB iron staining in the RAM of WT, *stop1* and *almt1* grown under -Pi and malate treatment (1 mM; -Pi/+Malate) conditions 5 dag. Scale bar equals 100 μm.

### Fe accumulation is absent in the RAM of Pi-deprived *stop1* and *almt1* seedlings and can be rescued by malate treatment

Accumulation of Fe in the apoplast of cells in the RAM is required to activate the primary root response to low Pi availability (20). As carboxylate-iron complexes have been reported to participate in iron transport and acquisition in plants (30-32) and malate efflux is affected in *stop1* and *almt1* mutants (21, 22), we explored whether malate exudation plays a role in the Fe accumulation mechanism that is required to trigger primary root growth inhibition in response to low Pi availability. First, we tested whether malate is required for the accumulation of Fe in the apoplast of RAM cells using Perls-DAB histochemical Fe staining, which allows the detection of changes in labile Fe^3+^ (20), on the root tips of WT, *stop1* and *almt1* in low Pi media with or without 1 mM malate (Figure 4D and Supplementary Figure 3). In Pi-deprived seedlings not exposed to malate, Fe staining was clearly observed in the roots of WT seedlings while *stop1* and *almt1* seedlings showed a much lower Fe accumulation (Figure 4D). In Pi-deprived WT seedlings treated with 1 mM malate, Fe staining was still clearly visible but in a more defined zone of the root apex, which included the QC. In contrast to *stop1* and *almt1* seedlings grown in -Pi media lacking malate, those treated with 1 mM of this organic acid showed a very similar pattern to that observed for the WT under the same conditions. Although malate-treated Pi-deprived *stop1* seedlings show a clear Fe staining, accumulation of Fe^3+^ in the RAM was apparently lower around the QC than that observed for the WT and *almt1* seedlings treated with malate (Figure 4C). We did not observe significant differences in the patterns of Fe staining between the root tips of WT, *stop1* and *almt1* seedlings under +Pi conditions treated with 1 mM malate (Supplementary Figure 3).

Citrate, as well as malate, is an organic acid that is released by plant roots in response to low-Pi availability (33). Since organic acids are naturally occurring metal chelating agents, and if the malate chelating-effect is responsible for the primary root growth inhibition in -Pi media, we wanted to test whether citrate treatment of Pi-deprived seedlings (1 mM citrate) could also phenocopy the short-root phenotype in Pi-deprived *stop1* and *almt1* seedlings. In -Pi media, we observed that citrate treatment slightly reduced primary root elongation of *stop1* seedlings (10%) (Supplementary Figure 4A-B) and had no significant effect in the root growth of Pi-deprived *almt1* seedlings (Supplementary Figure 4A-B). Interestingly, citrate treatment of Pi-deprived WT seedlings resulted in an average 2.5 fold increase in root length compared to that observed for WT seedlings grown in low-Pi media in the absence of citrate (Supplementary Figure 4A-B). These results suggest a specific role of malate in primary root growth inhibition by promoting the accumulation of Fe in the apoplast of root cells in the meristematic area.

As malate is capable of inducing root growth inhibition and meristem exhaustion in *almt1* seedlings, linked to an effect on Fe-accumulation in the RAM of *almt1* seedlings, we hypothesized that malate has a chelating effect on Fe that contributes to its accumulation in the root tip. To test our hypothesis, we performed molecular dynamic calculations to simulate the effect of malate on the aggregation of metallic ions such as Fe^2+^, Fe^3+^ and Al^3+^. We built 4 different simulation sets using ascending malate:metal molecular ratios, starting from 0:120 to 120:120 (Supplementary Figure 5). We observed non-bonded interactions between malate and Fe^2+^ ions but the interactions did not induce Fe^2+^ aggregation in any of the simulated systems (Supplementary Figure 5). In the case of the malate and Fe^3+^ system we observed non-bonded interactions and the formation of large malate-Fe^3+^ aggregates in all ratios tested with an increasing size of aggregates when an equimolar concentration of malate and Fe^3+^ was used (Supplementary Figure 5). Metals did not aggregate when malate was not included in the simulation set (Supplementary Figure 5). These results suggest that malate can form large aggregates with Fe^3+^ and Al^3+^ but not with Fe^2+^, which could be relevant for the activation of the *Arabidopsis* primary root response to low Pi.

### Differential expression analysis revealed a preferential loss of local transcriptional responses to Pi starvation in *stop1* and *almt1* root tips

The root tip plays a fundamental role in the ability of the root system to sense and respond to Pi starvation (16). Therefore, to understand the role of *STOP1* and *ALMT1* in the local and systemic responses of the *Arabidopsis* root to low phosphate, we performed a whole transcriptome sequencing (RNA-seq) analysis of gene expression in root tips from WT, *stop1* and *almt1* seedlings grown under +Pi and -Pi conditions (Figure 5). We performed pairwise comparisons of transcript abundances between -Pi and +Pi conditions to determine differentially expressed genes (−1.5 < logFC < 1.5; FDR<.05) in response to Pi deficiency in WT, *stop1* and *almt1* root tips (Figure 5A). A total of 1488 genes were found to be responsive to low phosphate in the WT (819 up; 669 down), while only 569 in *stop1* (294 up; 275 down) and 463 in *almt1* (224 up; 239 down) (Figure 5A). To identify the biological processes whose expression is misregulated in the root apex of *stop1* and *almt1* in response to Pi availability, we performed a Gene Ontology (GO) enrichment analysis (Figure 5). First, we performed a GO clusterization of all the over-represented categories that included genes that belong to the same biological process in the root tips of WT seedlings (Figure 5B). We found seven clusters that were named after the most significantly over-represented category of the cluster and included the cellular response to phosphate starvation (GO:0016036), secondary metabolism (GO:0019748), macromolecule metabolic process (GO:0044260), cell wall organization (GO:0071555) and systemic acquired resistance (GO:0009627). A full list of the GO categories that were enriched in the WT is included in Supplementary File 1. We then performed an analysis of the percentage of genes belonging to each cluster that were differentially expressed in *stop1* and *almt1* relative to the WT (Figure 5C). Using such an approach we determined that, overall, the percentage of genes that were activated in response to low phosphate in the root tip of the WT and were misregulated in *stop1* and *almt1* ranged between 40% to 95% (Figure 5C). The most affected biological process was “cell wall organization” as evidenced by the reduced number of transcripts from the cluster that were activated in response to low Pi in *stop1* and *almt1* (41 WT; 4 *stop1;* 2 *almt1*). Of the 46 differentially expressed genes included in the “response to phosphate starvation” that were regulated in WT root tips, 46% and 40% (21 *stop1;* 18 *almt1*) were differentially expressed in the root tips of *stop1* and *almt1*, respectively.

We found that the expression of *SPX1* and *SPX2*, two key genes in the regulation of Pi-responsive genes that are systemically induced (34), were normally induced in the root tips of *stop1* and *almt1* mutants (Supplementary File 1). Therefore, we examined the transcription levels of genes that are known targets of the PHR1/SPX systemic regulatory node (6). Of the 94 PHR1 direct targets that we found induced in the root tips of WT plants, we found that 60% (63 *stop1;* 57 *almt1*) were also induced in *stop1* and *almt1* in response to low Pi conditions (Figure 5D). Since the activation of genes in the “cell wall organization” cluster is one of the most affected in *stop1* and *almt1* and its transcriptional regulation has been recently linked to the local response to low Pi availability (35), our results suggest that the local response to low Pi is largely lost in *stop1* and *almt1* and that the systemic response to low Pi is significantly less affected than the local response in these two mutants. To confirm that this was indeed the case, we performed a comparison of genes that had been previously defined to participate in local and systemic responses to low Pi (5) with those that were not activated in *stop1* and *almt1* (Figure 5D). We found that among the 79 systemic and 147 local genes that were differentially expressed in the WT in response to Pi deficiency under our experimental conditions, over 60% of the systemically regulated genes remained responsive in *stop1* (52) and *almt1* (51), while less than 28 *%* of locally regulated genes (40 *stop1;* 24 *almt1)* remained responsive to Pi starvation in the root tips of the mutants (Figure 5D). These results confirm that *STOP1* and *ALMT1* have a key role in regulating the expression of genes in the local response to Pi deficiency. Nonetheless, *STOP1* and *ALMT1* also seem to have a significant effect on a group of genes that are systemically induced by low Pi availability.

**Figure 5.**
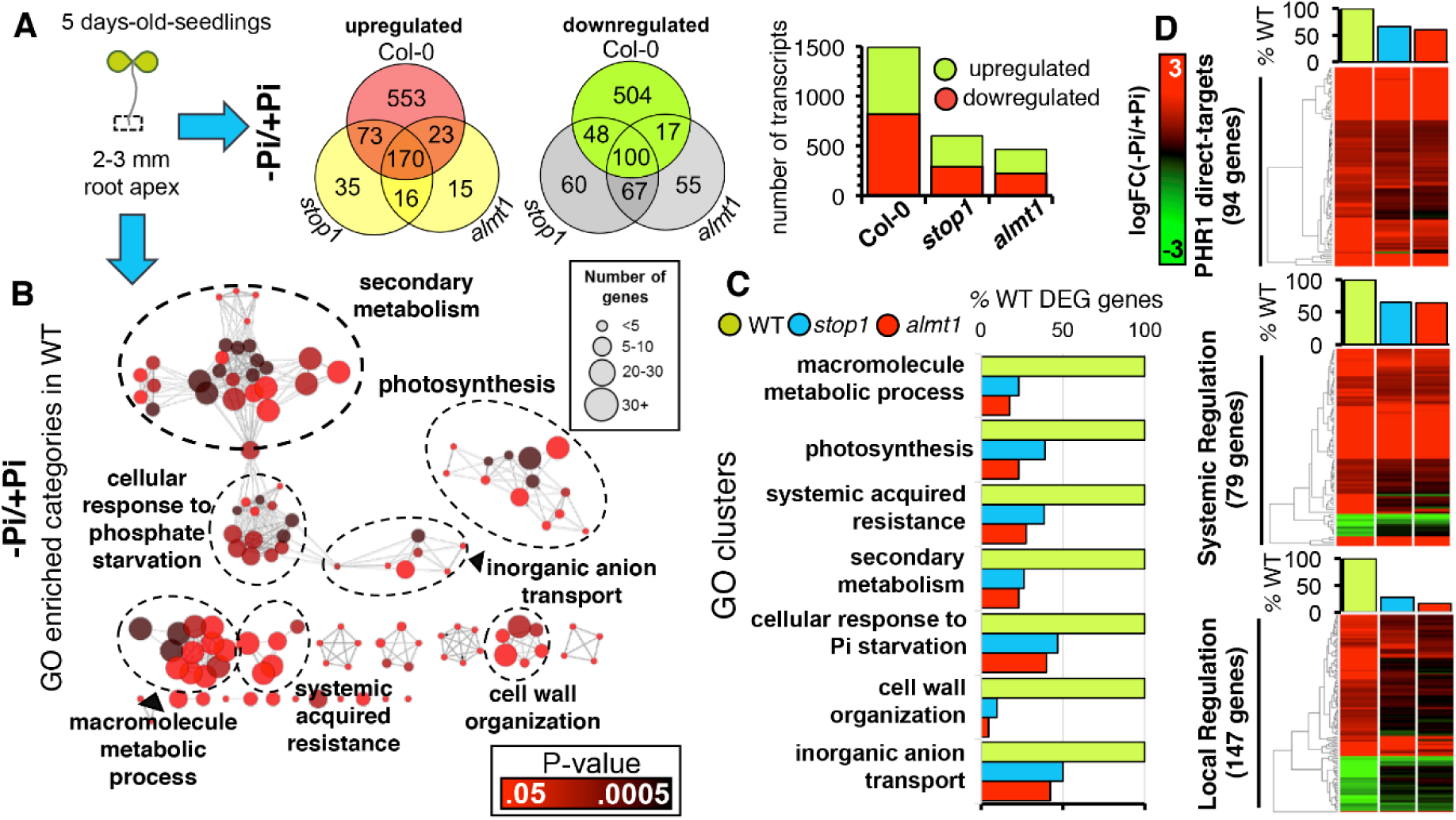
Differential expression profiling of WT, *stop1* and *almt1* revealed a loss of transcriptional response to Pi starvation in the root apex of *stop1* and *almt1*. (A) Venn diagram of differentially expressed genes (DEG) that are up- and down-regulated (−1.5 > logFC(-Pi/Pi) > 1.5; FDR < .05) in the root tip of WT, *stop1* and *almt1* seedlings in response to -Pi conditions. A bar-graph illustrating the number of upregulated and downregulated transcripts in WT, *stop1* and *almt1* is presented. (B) Gene Ontology (GO) enrichment analysis of overrepresented categories that are activated in the root apex of Pi-deprived WT seedlings. Each circle corresponds to a significantly enriched GO category (P-value <.05; hypergeometric test; Benjamini-Hochberg correction). Color code resembles P-value and size resembles the number of genes that are associated to that respective GO category. GO categories that share genes are connected and clustered by the biological process that corresponds to the most significantly enriched category of the cluster. (C) Analysis of DEG by cluster in the root tip of *stop1* and *almt1*. The number of genes that belong to each GO cluster and are differentially expressed in *stop1* and *almt1* is presented as a percentage of the number of genes that are differentially expressed in WT (% DEG of WT). (D) Transcriptomic analysis of locally and systemically regulated genes in the root apex of WT, *stop1* and *almt1* revealed a key role of *STOP1* and *ALMT1* in the local response to Pi starvation in *Arabidopsis*. WT, *stop1* and *almt1* logFC values of genes that are differentially expressed in the root tips of WT seedlings (-1.5 > logFC(-Pi/Pi) > 1.5; FDR < .05) in response to -Pi conditions and were reported as PHR1-direct targets (6) and of those that were classified as part of the local or systemic transcriptional response as reported by (5) is represented in a heatmap, respectively. Genes that are differentially expressed in WT root tips and their expression levels in WT, *stop1* and *almt1* seedlings grown under -Pi conditions are illustrated according to the key.

### Malate treatment rescued the expression of local-response genes encoding apoplast-located proteins

As we observed that malate treatment rescued iron accumulation in the RAM and the long root phenotype of *stop1* and *almt1* mutants, we sought to identify the subset of genes that regulate primary root growth inhibition and whose expression under low Pi conditions is re-activated by malate treatment in *stop1* and *almt1* seedlings. To this end, we isolated mRNA from the root tips of Pi-deprived *stop1* and *almt1* mutants that were treated with malate (1 mM) and carried out RNA-seq analysis (Figure 6). Our rationale was that the common set of genes that are differentially expressed in the root tips of Pi-deprived seedlings that have a short-root phenotype (WT, stop1+M, almt1+M) and that are not differentially expressed (induced or repressed) in the root tips of Pi-deprived *stop1* and *almt1* seedlings, which have a long root phenotype in -Pi media, are linked to the malate-dependent mechanism that triggers primary root growth inhibition under Pi deprivation conditions. A common set of 210 differentially expressed genes (63 upregulated; 147 downregulated) was found between the genotypes/treatment that induce a short root phenotype under -Pi conditions and that are not differentially expressed in Pi-deprived seedlings with a long root phenotype (Figure 6A). Among the genes whose expression was rescued by malate treatment, we found several peroxidase family genes (*PEROXIDASE2, PEROXIDASE37, AT3G01190, PEROXIDASE4*) which are closely related to the control of ROS homeostasis (36). A full list of the genes and description is included in Supplementary File 1. Interestingly, using SUBA, a subcellular prediction tool (Tanz et al. 2013), we found that 30% of proteins encoded by genes whose responsiveness to low Pi is restored by malate treatment in *stop1* and *almt1* Pi-deprived seedlings are targeted to the apoplast or extracellular region (Figure 6B), confirming a previous study in which a major role of the apoplast in the root response to Pi starvation was highlighted (35). Furthermore, an additional 16% of genes are targeted to the plasma membrane (Figure 6B).

**Figure 6.**
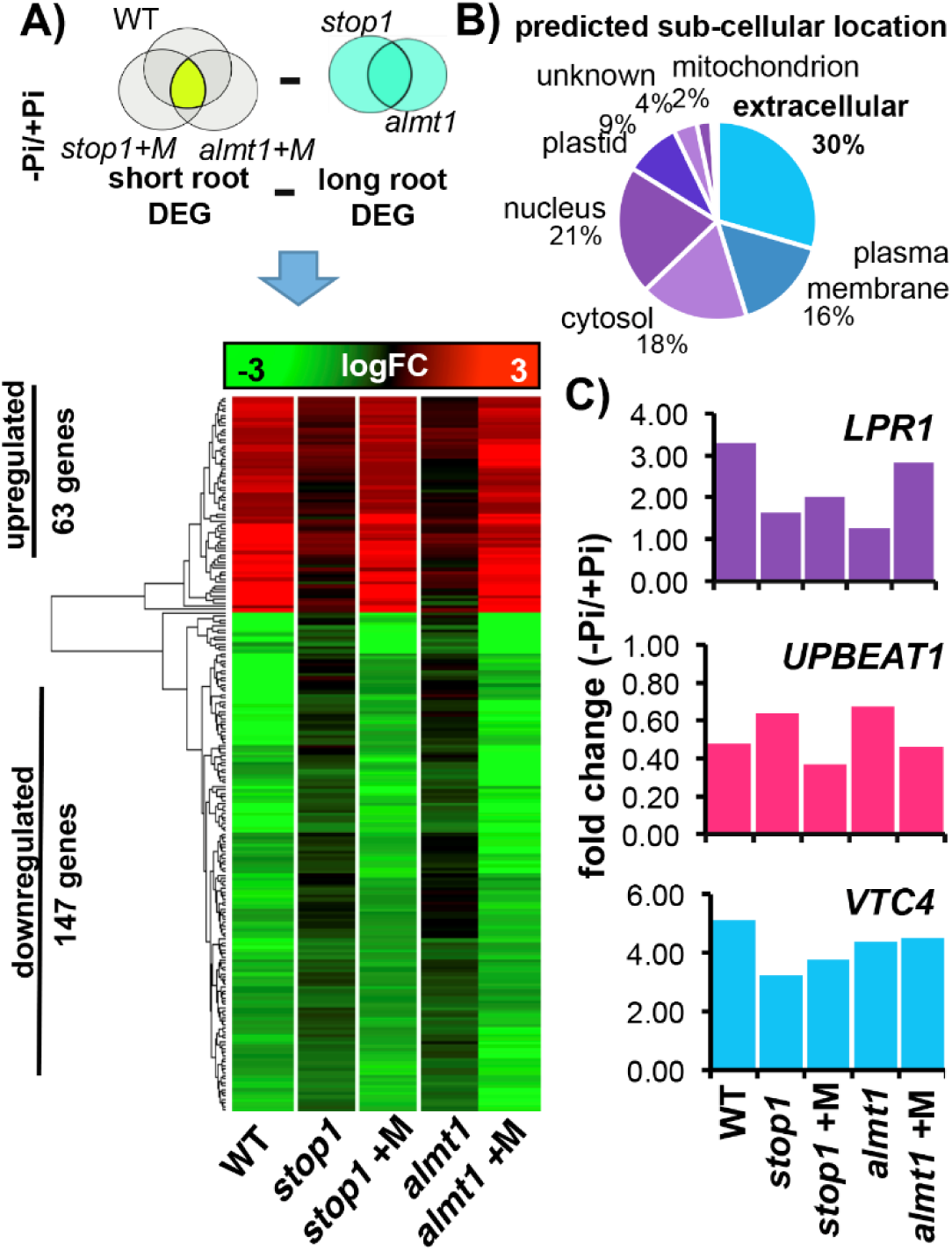
Malate treatment rescues the expression of transcripts whose products are targeted to the extracellular region. (A) Schematic of the Venn diagram analysis of common differentially expressed genes (DEG; −1.5 > logFC(-Pi/Pi) > 1.5; FDR < .05) in short-root phenotypes (WTՈstop1Ոalmt1) minus the DEG in long-root phenotypes (*stop1Սalmt1*) under Pi deficiency conditions, that was performed to determine the genes whose expression is linked to short-root phenotype and is rescued by malate treatment in *stop1* and *almt1* seedlings. A heatmap of the logFC values in WT, *stop1/+M* and *almt1/+M* of the determined gene set is presented. (B) Predicted subcellular location of the DEG whose expression is restored by malate treatment. (C) *LPR1*, *UPBEAT1* and *VTC4* expression levels are presented in fold change (FC; FDR<.05) as revealed by our transcriptomic studies in the root apex of WT, *stop1* and *almt1* seedlings under -Pi conditions and -Pi and malate treatment (*almt1*+M, *stop1*+M) conditions.

### LPR1 acts downstream of STOP1 and ALMT1

In *Arabidopsis, LPR1* has been reported to mediate the oxidation of Fe^2+^ to Fe^3+^ in the apoplast, which was correlated with ROS generation that triggers callose deposition in the root apex and disrupts SHR transport which ultimately induces determinate primary root growth in response to Pi deficiency conditions (20). As both Fe^3+^ accumulation and the regulation of peroxidases are lost and rescued by malate treatment in *stop1* and *almt1* mutants, using our RNASeq data, we analyzed if malate treatment had the same effect on the *LPR1* expression in response to low Pi conditions in the root apex of *stop1* and *almt1* (Figure 6C). The expression of *LPR1* was found to be enhanced in the root apex of WT seedlings (3.29 fold) exposed to low-Pi, induction that was significantly lower in *stop1* (1.64 fold) and *almt1* (1.25 fold) Pi-deprived seedlings (Figure 6C). Our data revealed that, indeed, malate treatment increased *LPR1* transcript levels in Pi-deprived *stop1* seedlings from 1.5 to 2.0-fold and in *almt1* seedlings from 1.2 to 2.7-fold (Figure 6C). The higher increase in *LPR1* expression induced by malate treatment in *almt1* than *stop1* correlates with the capacity of the treatment to better restore the primary root response to low in *almt1* than *stop1* (Figure 6C). To test whether the effect of malate was dependent or independent of LPR1, we studied the effect of malate treatment on the primary root elongation of *lpr1* seedlings grown in media lacking Pi. We observed that malate treatment did not rescue the *lpr1* mutant phenotype (Supplementary Figure 6), suggesting that the effect of malate to trigger the determinate root developmental program in response to Pi deficiency requires the presence of a functional LPR1 protein.

Since alterations in the ROS balance in the RAM of *Arabidopsis* are linked to the meristem exhaustion process observed in Pi-deprived seedlings (37) and *LPR1* is essential for the low-Pi dependent ROS signaling that takes place in the primary root of *Arabidopsis*, we decided to analyze the transcript levels of *UPBEAT1* (*UPB1;* Figure 6C); the only transcription factor known to module ROS balance and control the transition from cell proliferation to cell differentiation in the RAM by modulating the transcription of peroxidase genes (38). We observed that *UPBEAT1* is down-regulated 0.48-fold in response to low Pi in the root apex of WT seedlings and that it is downregulated to a lower degree in *stop1* (0.64 FC) and *almt1* (0.67 FC). Malate treatment restored the downregulation of *UPB1* in Pi-deprived *almt1* (0.46 FC) and *stop1* (0.37 FC) seedlings to WT levels (Figure 6C). Since LPR1 and UPB1 seem to be involved in modulating ROS balance in the RAM and their expression is misregulated in *stop1* and *almt1*, we identified peroxidase genes which have been related with ROS homeostasis in the root (36) and analyzed their expression levels in WT, *stop1* and *almt1* seedlings. We found that 18 peroxidase genes *(PRXS)* were transcriptionally regulated by low Pi in the root apex of WT seedlings of which 13 *PRXS* were not responsive to low Pi in *stop1* and *almt1* (Supplementary File 1). Interestingly, the responsiveness of 11 peroxidase genes *(PEROXIDASE52, AT4G08780, AT4G08780, PEROXIDASE2, AT5G06730, AT5G39580, PEROXIDASE37, AT5G15180, PEROXIDASE4, AT2G39040, AT3G01190)* was rescued by malate treatment in Pi-deprived *stop1* and *almt1* seedlings (Supplementary Figure 7). Alterations of ROS balance have been related to callose deposition in the primary root of *Arabidopsis* (20). Given that the transcription of ROS-related genes *(UPB1, LPR1 and PRXS)* is disrupted in *stop1* and *almt1* under low Pi conditions, we analyzed the expression values of genes coding for callose synthases *(CALS)*. We found that the expression of 4 genes coding for *CALS (CALLOSESYNTHASE7, CALLOSESYNTHASE9, GLUCANSYNTHASELIKE4, GLUCANSYNTHASELIKE5)* was induced (1.1 logFC; FDR<.05) in the root apex of WT seedlings and that it was induced to a lesser extent (logFC<0.6) in *stop1* and *almt1* seedlings. It was observed that malate treatment also rescued the expression of these 4 *CALS* in Pi-deprived *stop1* and *almt1* (Supplementary Figure 7). Our results suggested that malate efflux is required for the ROS signaling cascade that has been reported to be affected in low-Pi insensitive mutants (20, 37).

Given that Fe tends to its higher oxidation state (Fe^3+^) in natural environments and the iron uptake genes *IRON REGULATED TRANSPORTER 1* (*IRT1*) and *FERRIC REDUCTION OXIDASE 2* (*FRO2*) have been reported to be repressed in response to Pi deficiency conditions (3, 5, 39) we asked whether an alternative Fe^3+^ reduction mechanism could provide Fe^2+^ to LPR1 to initiate the proposed ROS signaling cascade that is induced in the RAM in response to low Pi. An ascorbate dependent Fe^2+^ reduction mechanism has been reported recently (31), in which *VITAMINC4* (*VTC4*), a gene encoding a protein with dual myo-inositol-monophosphatase and ascorbate synthase activity (40), could play a central role. We found that *VTC4* is induced in the root apex of WT (5 FC), *almt1* (4.3 FC) and *stop1* (3.22 FC) seedlings (Figure 6C). Interestingly, *VTC4* belongs to the set of genes that are direct targets of PHR1 (6), providing a potential link between local and systemic signaling in the primary root response to low Pi and the crosstalk between Fe and P in the Pi deficiency response.

## Discussion

LPR1 has been proposed to promote the accumulation of Fe in the apoplast of cells in the RAM, which in turn triggers an accumulation of callose that alters symplastic transport causing meristem differentiation (20). However, the precise mechanism by which Fe accumulates in the apoplast of RAM cells remained to be determined. Here we show that *STOP1* and *ALMT1* participate in the mechanism that triggers RAM exhaustion in response low Pi availability by mediating the accumulation of Fe^3+^ in the apoplast of RAM cells. *STOP1* and *ALMT1* were originally described as genes responsible of the malate efflux that protects the *Arabidopsis* root from Al^3+^ toxicity (21, 22). *AtSTOP1* is constitutively expressed in *Arabidopsis*, indicating that its involvement in the Al-dependent induction of gene expression must involve posttranslational processes of modification or the direct binding of Al^+3^. The finding that mutations in *STOP1* and *ALMT1* lead to long root phenotypes in Pi-deprived seedlings suggests that malate excretion is also involved in the process of meristem exhaustion in response to low Pi availability. We found that *STOP1* is expressed in the RAM in a Pi-independent fashion and that *ALMT1* is expressed in the RAM but only in seedlings grown in media with low Pi concentrations. Although the expression domains of *STOP1* and *ALMT1* do not completely overlap, we found that the expression directed by the *ALMT1* promoter is completely dependent on an intact copy of *STOP1* (Supplementary Figure 2). The apparent inconsistency between the role of an activator with its target gene and the differences in patterns of expression of *STOP1* and *ALMT1* could be explained by the recent report that *STOP1* mRNA is cell-to-cell mobile (42).

Pi and Fe availability have been shown to coordinately regulate RAM maintenance and primary root growth in vitro (11, 17, 20). Our results corroborate that, in *Arabidopsis* WT seedlings, Fe availability (Supplementary Figure 8) in the medium is required for RAM-exhaustion in media with a low Pi concentration and that Fe accumulation in the RAM is associated with the process of RAM- exhaustion (Figure 4). Apoplastic iron accumulation in the RAM was reported to be essential for primary root growth inhibition in response to -Pi conditions (20), however, the mechanism for iron accumulation in the root remained to be determined. We found that Fe failed to accumulate in the root apex of *stop1* and *almt1* seedlings grown in Pi-deficient media and that the treatment of *stop1* and *almt1* seedlings with malate restores both Fe accumulation in the RAM and the inhibition of primary root growth in Pi-deprived seedlings (Figure 4). These data show that malate secretion is necessary and sufficient for iron accumulation in the RAM and to trigger cell differentiation in the RAM that is responsible for the meristem exhaustion process induced by Pi deficiency. We propose that such mechanism of iron accumulation happens in the apoplast as ALMT1 is reported to be a malate efflux protein (22) and thus, exogenous malate, which probably diffuses through the apoplast, can rescue iron accumulation in *almt1* and *stop1*. Malate supplementation was found to fully rescue the short-root phenotype of *almt1* while it only partially restored primary root growth inhibition in *stop1*, suggesting that, in addition to *ALMT1, STOP1* regulates the expression of other genes whose activation by low Pi is required for full meristem exhaustion.

Molecular dynamic simulation of Fe^3+^ and Fe^2+^ interaction shows that malate can form large complexes with Fe^+^ but not Fe^+2^ (Supplementary Figure 5). This data suggests that malate promotes the accumulation of Fe^+3^ in the apoplast by forming these large molecular weight complexes, which by a still largely unknown mechanism that correlates with ROS generation (20), activate the processes required for meristem exhaustion. *lpr1* seedlings also show a long root phenotype in low Pi media, suggesting that the ferroxidase activity of LPR1 is required to trigger cell differentiation during the primary root meristem exhaustion process triggered by Pi-deprivation. We found that the long root phenotype of *lpr1* in low-Pi media cannot be rescued by malate treatment, suggesting that *LPR1* acts downstream of *STOP1* and *ALMT1* and that most probably is required to activate the Fe-mediated mechanism involved in the process of RAM exhaustion observed in Pi-deprived seedlings. Therefore, cell-wall-localized LPR1 ferroxidase activity which catalyzes Fe^+2^ to Fe^+3^ conversion (20), could act synergistically with malate efflux in the accumulation of Fe^3+^in the apoplast of RAM cells. LPR1-dependent Fe^3+^ production in the apoplast could trigger ROS production by initiating a Fe redox cycle as previously proposed (43). Either ferric-chelate reductase oxidase activity, which reduces apoplast-diffusible Fe^3+^chelates, or effluxed ascorbate (31)could reduce the Fe^3+^ produced by LPR1 to a redox-active Fe^2+^ to complete a cycle thereby triggering root cell differentiation. Supporting this notion, *LPR1* overexpression causes ectopic Fe^3+^ and ROS generation in Pi-deprived seedlings (20). Our data suggest the existence of a *STOP1, ALMT1* and *LPR1* coordinated redox mechanism that involves Fe^+3^ deposition in the apoplast of RAM cells of seedlings exposed to low Pi. *ALMT1* is transcriptionally upregulated in a similar fashion in WT and *lpr1* mutants (35), confirming that LPR1 acts downstream of the STOP1/ALMT1 low Pi regulatory node.

Expression of up to 80% of the genes involved in the local response to low Pi was affected in *stop1* and *almt1*, whereas less than 40% of Pi systemically responsive genes were affected in the same mutants (Figure 5D). These results show that, in addition to controlling the primary root developmental response to low Pi, *STOP1* and *ALMT1* play an important role in modulating the transcription of other genes involved in the local response to Pi starvation. However, the reduction in both local and systemic responses in *stop1* and *almt1* points to a cross-talk between the signaling pathways that regulates the transcriptional activation or repression of the systemic and local responses to low Pi in the root apex. However, we cannot rule out the possibility that the internal concentration of Pi could be higher in the root tip of *stop1* and *almt1* than the WT, which could lead to a downregulation of the systemic response, as has been observed in plants grown in low Pi and low Fe conditions (17). Further experiments regarding a possible STOP1 and ALMT1 interaction directly with Pi, PHR1 or SPX-domain proteins could shed light on the existence of a coordinated response to external and internal Pi levels in the root apex. The observation that PHR1 activates the expression of the ascorbate synthase VTC4 under Pi deficiency conditions together with the recent report that ascorbate efflux contributes to Fe^3+^ reduction (31), support the notion that the redox cycle that generates ROS and triggers RAM exhaustion could be controlled by both local and systemic responses to Pi starvation.

Our transcriptomic analysis revealed that *LPR1* is responsive to Pi-deprivation in the root tips of WT plants (Figure 6C) and that this response is significantly reduced in the root tips of *stop1* and *almt1* seedlings (Figure 6C). These results suggest that a threshold level of *LPR1* is required to activate meristem exhaustion in Pi-deprived seedlings. This notion is supported by the observation that the treatment with malate that reverts the long root phenotype of Pi-deprived *stop1* and *almt1* seedlings (Figure 4A) also leads to an increase in *LPR1* transcript levels (Figure 6C). Moreover, *Arabidopsis* accessions with higher *LPR1* transcription levels have shorter primary roots under low Pi conditions (16) and we observed that, in the case of malate-treated Pi-deprived *stop1* and *almt1* seedlings, a higher *LPR1* expression in *almt1* than in *stop1* correlated with a shorter primary root in *almt1* than *stop1* (Figure 4B, 6C). A recent report on the interplay between the transcriptional activation of genes coding for extracellular enzymes and iron-redistribution in the apoplast in response to Pi-deprivation highlighted the role of the apoplast in the Pi starvation response (35). Transcriptomic analysis of Pi-deprived *stop1* and *almt1* seedlings showed that malate treatment reactivates the low-Pi-responsiveness of genes that encode extracellular proteins involved in cell-wall modification and ROS homeostasis, such as peroxidases (36). Our transcriptomic results confirm a previously reported role of apoplastic peroxidases in the Pi starvation response (35), and highlight the role of malate secretion in the cell wall remodeling processes potentially involved in the changes of *Arabidopsis* root system architecture induced by low Pi availability. In this context, the finding that *UPBEAT1*, a transcription factor that modulates the transition from cell proliferation to cell differentiation in the RAM by repressing peroxidase genes (38), is repressed in response to Pi deficiency conditions and the fact its expression is altered in *stop1* and *almt1*, support the notion that ROS generation plays an important role in the root response to Pi deprivation. Further experiments regarding the specific pattern of ROS signaling in the RAM under Pi starvation conditions are required.

A model that summarizes what is known about the local response to Pi starvation and the proposed role of *STOP1* and *ALMT1* in the root response to Pi deprivation is presented in Figure 7. Under Pi deficiency conditions, expression of LPR1 is enhanced by a malate-dependent mechanism and PDR2 activity is inhibited facilitating LPR1 mobilization from the ER to the plasma membrane (20) where LPR1 ferroxidase activity catalyzes Fe^2+^ to Fe^3+^ conversion in the apoplast of RAM cells (Figure 7). The mechanisms by which LPR1 is transported from the ER to the extracellular region and how PDR2 activity is regulated by Pi-availability remain to be determined. *STOP1, a* constitutively expressed gene, up-regulates the expression of *ALMT1* in seedlings exposed to low Pi, thereby activating the excretion of malate. LPR1 ferroxidase activity in the plasma membrane of cells in the meristem and elongation zones of the primary root locally produces Fe^3+^ which forms large complexes with malate, leading to its accumulation. Expression of the ascorbate synthase VTC4 under Pi deficiency conditions is activated and the presence of ascorbate in the apoplast produces the Fe^2+^ required to complete a redox cycle that generates ROS. Expression of the *UPB1* repressor is reduced by a Fe/malate-dependent mechanism under low Pi conditions, which enhances the transcription of peroxidase genes. Enhanced transcription of local response peroxidase genes likely triggers ROS generation in the root of Pi-deprived seedlings as previously reported (37). ROS generation triggers callose deposition as previously reported (20, 44). This notion is supported by the observation that the expression of several *CALS* genes is enhanced in the root apex of WT but not in *stop1* and *almt1* seedlings (Supplementary Figure 7). Callose synthesis impairs symplastic transport by physically blocking plasmodematal pores, which reduces or inactivates the cell to cell movement of SHR (20). Since SHR cell to cell movement is required for stem cell niche maintenance, meristem exhaustion takes place in seedlings exposed to low Pi availability. However, since CLE-like peptide signaling is also required for stem cell niche maintenance in *Arabidopsis* (45) we cannot rule out that ROS or Fe^3+^ accumulation in the RAM could induce the transcription of CLE peptides that could also execute RAM exhaustion.

STOP1 controls *ALMT1* transcription and its expression is not regulated by Pi availability, suggesting that STOP1 is most likely involved in sensing external Pi-levels or an environmental cue that it is linked to low-Pi levels in the medium. Since STOP1 also controls low pH and Al^3+^ toxicity responses, it emerges as a possible master regulator/sensor that orchestrates the root responses to multiple environmental stresses.

**Figure 7.**
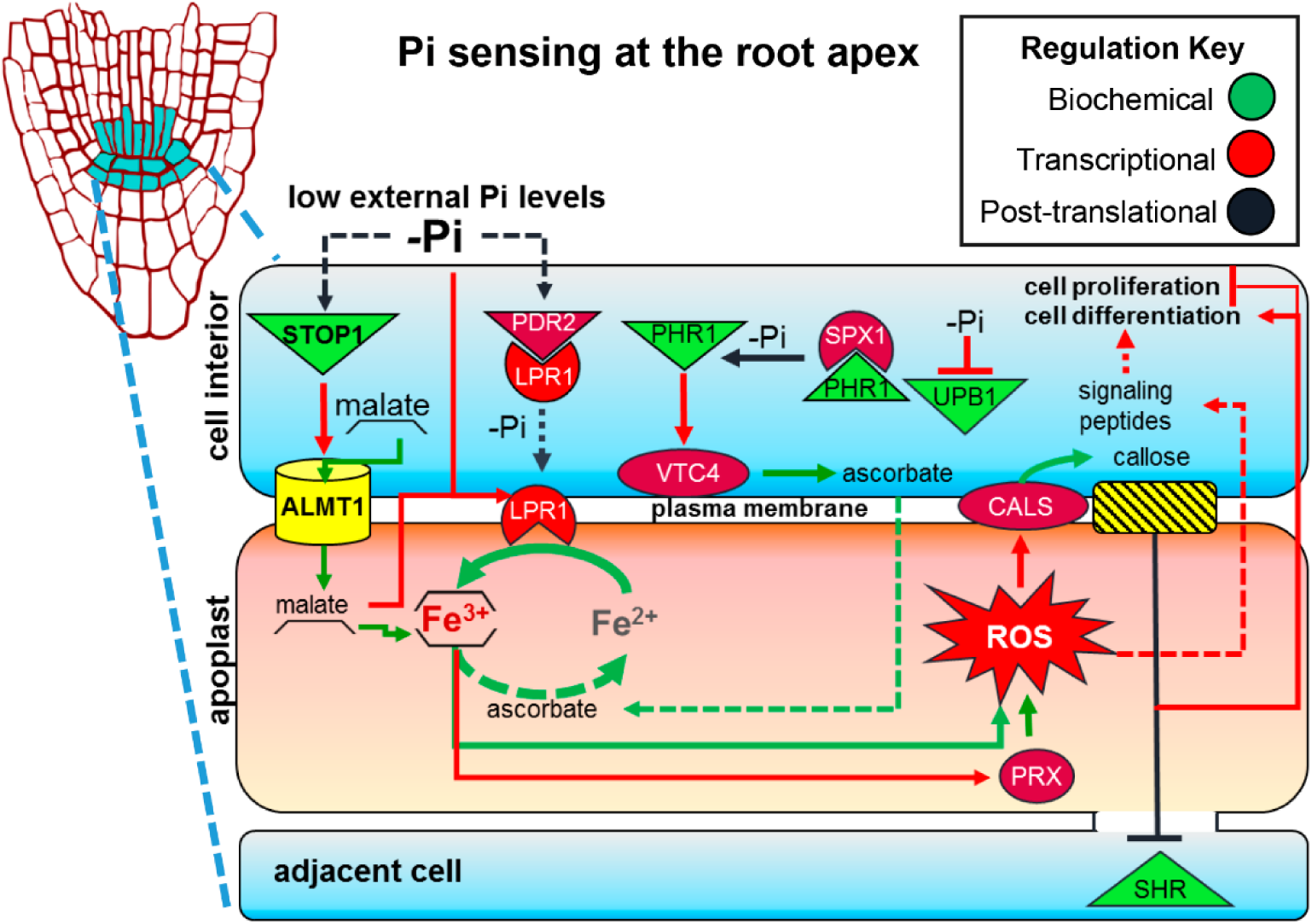
*STOP1* and *ALMT1* regulate the RAM response to low Pi conditions in *Arabidopsis*. In response to limiting Pi levels in the medium, *STOP1* induces *ALMT1* (Figure 3) which triggers malate efflux in the root apex. Low Pi levels also inhibit PDR2 negative regulation over LPR1 and induce, in a largely unknown mechanism, LPR1 transport to the plasma membrane. Malate contributes to the aggregation of Fe^3+^ ions in the apoplast and enhances the expression of *LPR1*, callose synthase genes *(CALS)* peroxidase genes *(PRX)* under Pi deficiency conditions. LPR1 and PRX activity generates reactive oxygen species (ROS) which enhance callose deposition by CALS. Callose deposition closes the symplastic channels of communication which ultimately impairs the transport of transcription factors, such as SHR, that are essential to maintain cell proliferation and organization in the RAM. Alternative factors induced by ROS, such as signaling peptides, could also induce cell differentiation in the RAM. A crosstalk between internal sensing, mediated by the PHR1/SPX1 module, and external Pi-sensing could be interconnected by ascorbate efflux into the apoplast, which can reduce Fe^3+^ to Fe^2+^ and could re-start the proposed redox cycle. Ascorbate can be produced by VTC4 whose gene is induced by PHR1 when internal Pi levels are limiting. Arrows represent relationships between the components. Dotted lines represent hypothetical relations, and the regulation key illustrates the type of evidence that has been provided for that relationship.

## Materials and Methods

### Plant material

*Arabidopsis thaliana* Col-0 accesion (CS70000) was used in this work. Stop1-SALK_114108 (N614108), almt1-SALK_009629 (N509629), and lpr1 (N516297) lines were provided by the NASC (European Arabidopsis Stock Center).

### Growing conditions

Seeds were surface sterilized and sowed in 1% agar, 0.1x MS medium as described by (10). 1mM KH_2_PO_4_ (high Pi; +Pi) or 10μM KH_2_PO_4_ (low Pi; -Pi) Pi concentrations were used. 1% (w/v) sucrose and 3.5mM MES was added. Fe-free medium was prepared as described (15), and 100μM ferrozine (SIGMA-82950) was added to reduce agar Fe-availability. Malate and citrate (SIGMA-M1000 and SIGMA-C0759, respectively) were added to medium before sterilization. Seedlings were grown in a Percival chamber at 22 °C, under 16/8 hrs photoperiod with >200 μmol^*^m-2^*^−1 luminous intensity.

### EMS mutagenesis

Over 3000 *Arabidopsis* seeds were surface sterilized with ethanol 95% (v/v) ethanol for 10min and 20% (v/v) bleach for 6 min. The sterilized seeds were left overnight at 4°C on a rocker in distilled water. Then seeds were incubated with 0.04% EMS (SIGMA-M0880) during in 100 mM sodium phosphate buffer, pH7. After 9 hours the seeds were washed ten times with distilled water to remove EMS residues. EMS mutagenized seeds (M1) were propagated under greenhouse conditions and F1 seeds (M2) were harvested and grown under -Pi conditions. Mutants that presented a long-root phenotype under in -Pi media were selected.

### Gene mapping

Plant mapping populations were built crossing *lpi5* and *lpi6* homozygous lines vs Col-0 (CS70000) respectively, according to the Mutmap protocol (26). Heterozygous F1 plants were self-fertilized for seed propagation. The F2 segregating individuals, were re-screened for WT (short root) and mutant (long root) phenotypes, respectively for each of the two crosses. Then, DNA was extracted from 100 plants for each phenotype, pooled and sequenced using Illumina Miseq technology (paired-end, 250 base-pair reads length and 50X coverage). Reads obtained were processed with fastQC and Trimmomatic (46) to improve their quality. Only paired-end reads were considered. Reads were mapped to the Col-0 reference genome (TAIR10) using BWA (47) and SAMtools (48). To identify and to evaluate specific variants related with the mutant phenotype (long root) a pipeline using GATK (https://software.broadinstitute.org/gatk/), VCFtools (49), SNPeff and SNPshif (27) was implemented. IGV was used to visualize the variants analyzed (50).

### Transcriptional reporter lines

Transgenic Col-0 plants harboring transcriptional gene fusions containing the *STOP1* and *ALMT1* promoter regions, respectively, fused to a double GFP-GUS reporter gene were produced using4900pb (primers: stop1Fw:GAACGACAAGATTACAAGTAGGTTC and stop1 Rv:GTTGCACAAATCGTCTTCAGTTTCC) and 2253 (primers: almt1 Fw:GGCAGATAAAGAGGCACTCGTG and almt1Rv:CTCTCTCACTTTCTCCATAACACC) intergenic regions of *STOP1* and *ALMT1*, respectively, were used to build the *proSTOP1::GUS-GFP* and *proALMT1::GUS-GFP* transcriptional reporter lines respectively. Intergenic region were cloned on the pKGWFS7 using the Gateway system. *Arabidopsis* plants were agro-infiltrated using the floral-dip method (51).

### Histochemical GUS staining

Histochemical GUS staining was performed as reported by (52). The stained roots were clarified following the protocol reported by (53). A representative root stained was chosen and photographed using Nomarski optics on a Leica DMR microscope.

### Histochemical iron staining

Perls iron staining and DAB intensification were carried out as described in (20) and analyzed using Nomarski optics on a Leica DMR microscope.

### Malate and Fe(II), Fe(III) and Al(III) Molecular Dynamics

Molecular dynamic (MD) calculations were carried out to obtain insight about the behavior between malate molecules and Fe(II), Fe(III) and Al(III) ions in explicit water solution. Non-bonded parameters for the metallic cations were taken from the 12-6-4 Lennard-Jones-type non-bonded model for divalent (54) and highly charged metal ions (55) respectively. These parameters were then adapted to the GROMOS 53a6 force field and integrated with GROMACS 5.0 as a user specified non-bonded potential employing tabulated interaction functions (56). The all-atom PDB optimized geometry structure and parameters for malate molecules were taken from the ATB web server (57). A total of nine systems were built with the malate-Fe(II), malate-Fe(III) and malate-Al(III) proportions of 0:120, 40:120, 80:120 and 120:120 within a cubic box with periodic boundary conditions. In all systems, the box was solvated with SPC/E (58) type water molecules and the steepest descents method was employed to minimize the energy. The temperature was set to 300 K and an equilibration phase of 100 ps in the canonical ensemble (NVT) was conducted using the V-rescale algorithm (59). Long-range electrostatics were calculated employing the PME method (60, 61) with a cutoff of 12 Å and the same cutoff was chosen for the van der Walls non-bonded interactions. All bond lengths were constrained with Linear Constraint Solver (LINCS; (62). A final production of 50 ns in the isothermal-isobaric ensemble (NPT) was conducted using the Parrinello-Rahman algorithm (63). Snapshots were stored after each 10 ps and the final MD trajectories were analyzed using the g_aggregate tool (64).

### Preparation of root tip mRNA-seq libraries

Total RNA was isolated from frozen root tip powder using TRIzol reagent (Invitrogen) according to the manufacturer’s instructions. Frozen root tip powder was obtained from root tip sections of approximately 2-3 mm length from approximately 500 individuals per treatment (5 dag). Non-strand specific mRNA-seq libraries were generated from 5 μg of total RNA and prepared using the TruSeq RNA Sample Prep kit (Illumina) according to the manufacturer’s instructions.

### RNA-seq High-throughput Sequencing and Data Analysis

High-throughput sequencing and data analysis were carried out as in (65). Briefly, quality assessment of the reads generated with the Illumina analysis pipeline (fastq format) was performed using FastQC (version 0.11.4) and processed using Trimmomatic (46) (version 0.35) to remove reads that contained adapter sequences and low quality reads. Single and paired-end clean reads were aligned to the Arabidopsis thaliana TAIR10 reference sequence using TopHat2 (66) (version 2.0.9). Raw counts per gene were estimated using HTseq (67) (version 0. 6.0). Data was normalized in edgeR (68) (version 3.12.0) using the trimmed mean of M values (TMM) method. Genes with ≥ 3 reads in total, across all samples, were included in the final analysis. Transcript abundance as represented by the normalized raw counts per gene was used to determine differential expression using the edgeR package. Analysis of GO enriched categories and clusterization into functional groups by biological process was performed using Cytoscape (69) (version 3.4) plugin ClueGO+CluePedia (70).

### qRT-PCR

Total RNA was isolated using TRIzol reagent (Invitrogen) according to the manufacturer’s instructions. Real-time PCR was performed with an Applied Biosystems 7500 real-time PCR system using SYBR Green detection chemistry (Applied Biosystems) and gene-specific primers. The relative expression levels were computed by the Ct method of relative quantification. Oligonucleotide primer sequences are available upon request.

## Acknowledgements

We thank G.S. Gillmor for his advice on EMS seed mutagenesis. We thank L.F. García-Ortega and O. Martínez for providing TransInfo: an algorithm to calculate the dispersion values for the normalization of transcriptomic data expression (unpublished). J.M.M. is indebted to Consejo Nacional de Ciencia y Tecnología (CONACyT) for a PhD fellowship. J.O.O.R is indebted to CONACyT for an MSc fellowship. This research was supported by CONACyT via general support to Laboratorio Nacional de Genómica para la Biodiversidad and by Howard Hughes Medical Institute Grant 4367 (to L.H.E).

## Competing Interest

The authors declare no competing interest.

Supplementary File 1 (.xlsx format) is deposited in the following link:https://www.dropbox.com/s/u5fv1wbrcmd1th6/Supplementary%20File%201.xlsx?dl=0

Genomic data generated in this study is accessible at www.ncbi.nlm.nih.gov/biosample/ Accession codes:

WT-*lp/5*: SAMN06013467; Mutant-*lp/5*: SAMN06013468; WT-*lp/6*: SAMN06013469;

Mutant-*lp/6*: SAMN06013470;

http://www.ncbi.nlm.nih.gov/biosample/6013467

http://www.ncbi.nlm.nih.gov/biosample/6013468

Expression data generated in this study is deposited in the Gene Expression Omnibus database (GSE90061) and can be accessed at:

https://www.ncbi.nlm.nih.gov/geo/query/acc.cgi?token=gfybucyuzbwrdyn&acc=GSE90061

## Supplementary Material

**Supplementary Table 1.** Segregation ratios of the F2 progeny seedlings obtained from *lpi5* X WT and *lpi6* X WT crosses.

| Cross | *F1 progeny* |  | *F2 progeny* |  |  |  |  |  |  |
| --- | --- | --- | --- | --- | --- | --- | --- | --- | --- |
|  | Phenotypes |  | Phenotype observed |  | Ration obtained |  | Ration tested |  | X<sup>2</sup>* |
|  | WT | mutant | WT | mutant | WT | mutant | WT | mutant |  |
| *lpi5* x *Col-0* | 179 | 0 | 490 | 170 | 2.97 : 1.03 |  | 3 : 1 |  | 0.2 |
| *lpi6* x *Col-0* | 148 | 0 | 577 | 182 | 3.04 : 0.96 |  |  |  | 0.42 |
| a95% |  |  |  |  |  |  |  |  | 3.841 |

\* With one degree of freedom and a critical value of 5% we fail to reject the hypothesis of a Mendelian ratio of segregation (3:1, WT:mutant) if the $\chi^2$ is smaller than 3.841.

**Supplementary Figure 1.**
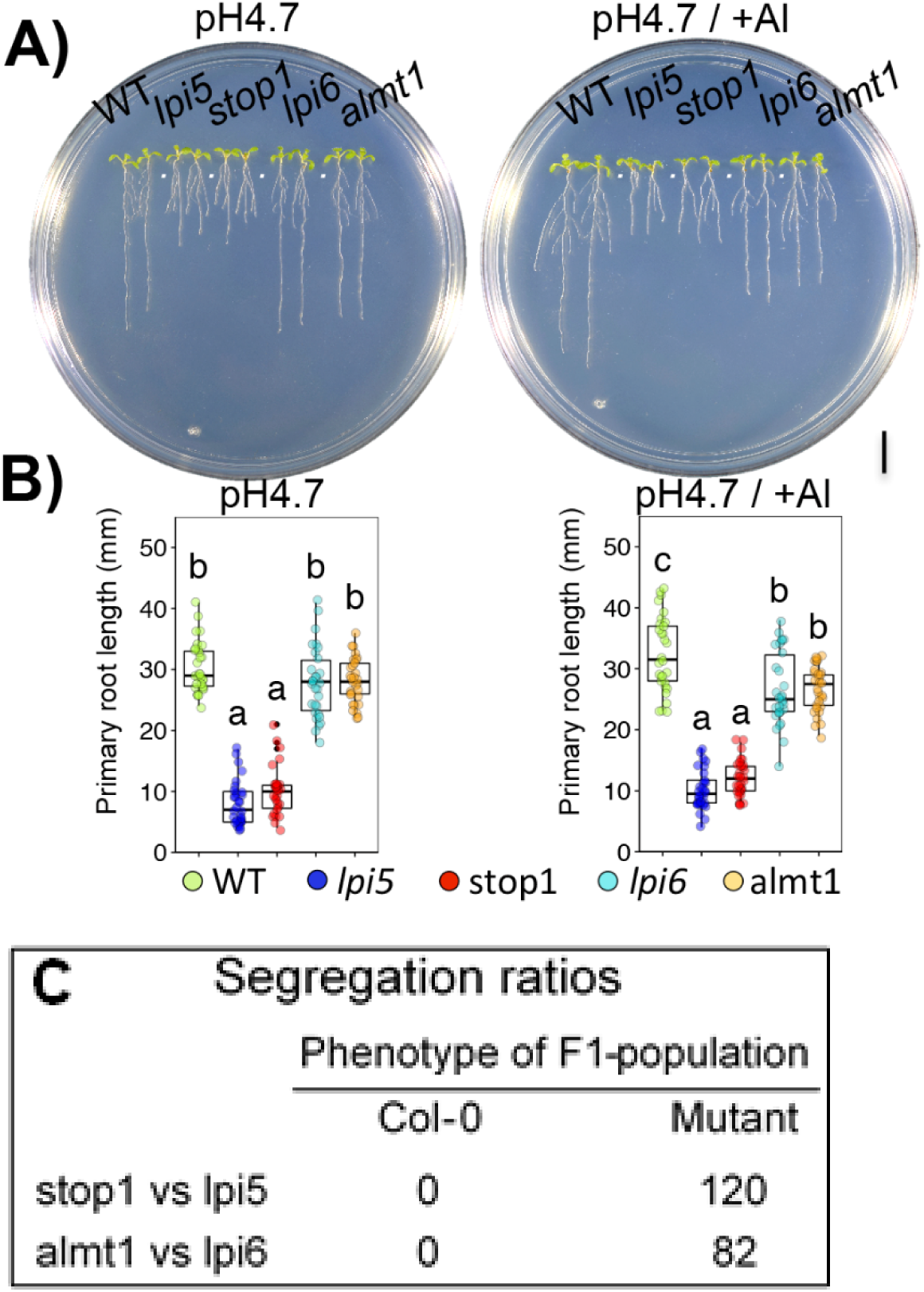
Primary root growth of *lpi5, stop1, lpi6* and *almt1* 10-day-old seedlings in response to low pH and aluminum toxicity. (A) Boxplot representation of the primary root length of WT, *lpi5, stop1, lpi6*, and *almt1* 10-day-old individuals (n=30) grown under low pH (4.6) (left) and aluminum toxicity (2 μM) (right) conditions. Dots represent individuals and genetic backgrounds are depicted by colors as described. Statistical groups were determined using a Tukey HSD test (P-value < .05) and are indicated by a letter. B) WT, *lpi5, stop1, lpi6*, and *almt1* 10-dag seedlings grown under low pH (4.6) (left) and aluminum toxicity (2 μM) (right) conditions. Scale bar equals 1 mm. C) Segregation ratios of F1 progeny seedlings (long root phenotype under -Pi conditions) obtained from *stop1* vs *lpi5* and *almt1* vs *lpi6* crosses. All F1 progeny seedlings from the *lpi5* x *salk_114108* and *lpi6* x *salk_009629* crosses were observed to have a mutant phenotype under -Pi conditions.

**Supplementary Figure 2.**
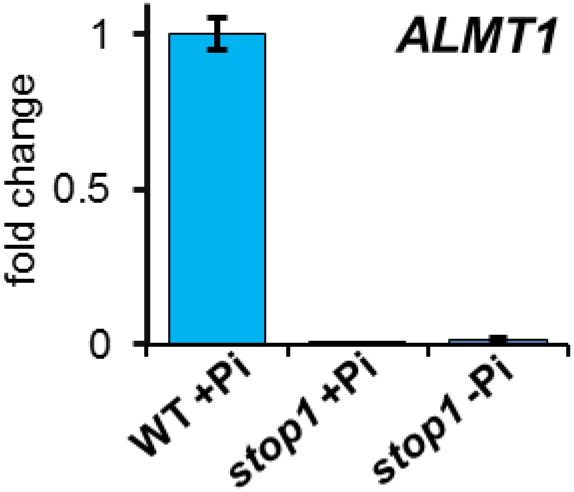
*STOP1* is essential for *ALMT1* expression in the root tip. qRT-PCR analysis of *ALMT1* expression in root apex of WT and *stop1* plants. Bars represent the mean logFC ±SEM of 2 biological replicates with 3 technical replicates. WT +Pi samples were used as calibrator values. *ACT2* and *UBQ10* were used as internal controls.

**Supplementary Figure 3.**
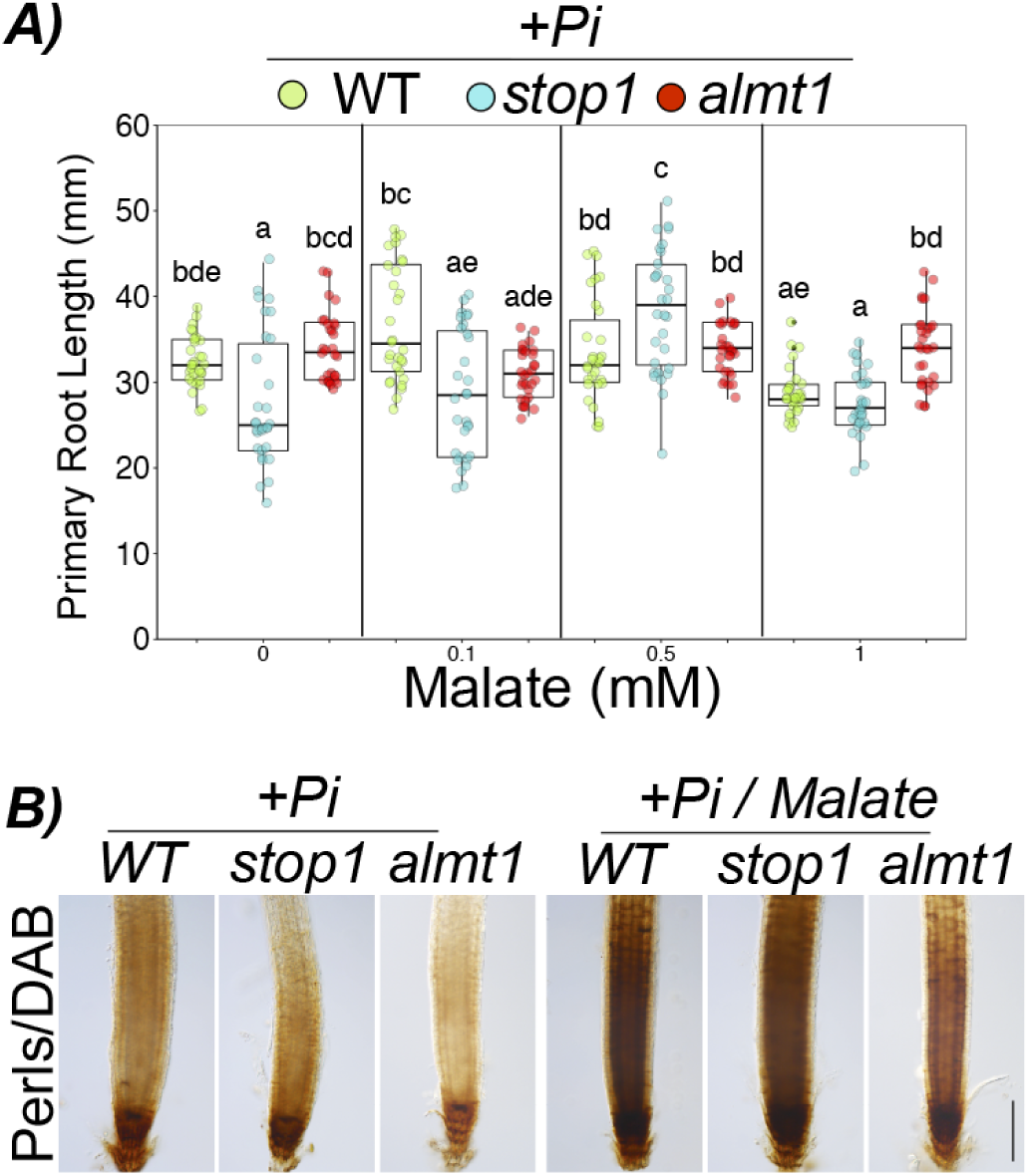
Malate effect on primary root growth and iron distribution in the root under high phosphate conditions. (A) Primary root length of 10 dag WT, *stop1* and *almt1* seedlings in response to increasing concentrations of malate supplemented to +Pi medium. (B) DAB-Perls iron staining of roots from 10 dag Col-0, *stop1* and *almt1* seedlings grown under +Pi and +Pi medium supplemented with 1 mM malate (+Pi+M).

**Supplementary Figure 4.**
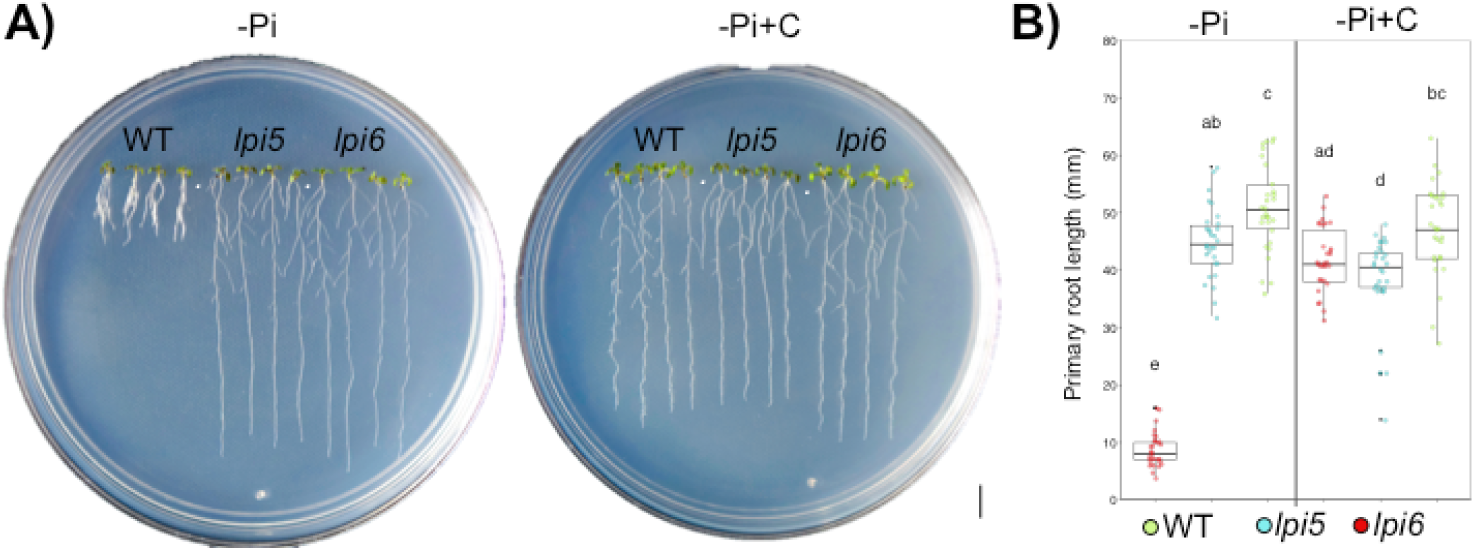
Citrate supplementation of low Pi medium did not rescue the long root phenotype of *stop1 and almt1* mutants. (A) Phenotypes of 10 dag WT, *stop1* and *almt1* seedlings grown in low phosphate medium (-Pi) and low phosphate medium supplemented with 1mM citrate (-Pi+C). Scale bar equals 1 mm. (B) Primary root length of 10 dag WT, *stop1* and *almt1* seedlings under -Pi and -Pi medium supplemented with 1mM citrate. Green, blue and red dots depict WT, *stop1* and *almt1* individuals (n=30 from 3 independent experiments), respectively. Statistical groups were determined using Tukey HSD test (P-value <.05) are indicated with letters.

**Supplementary Figure 5.**
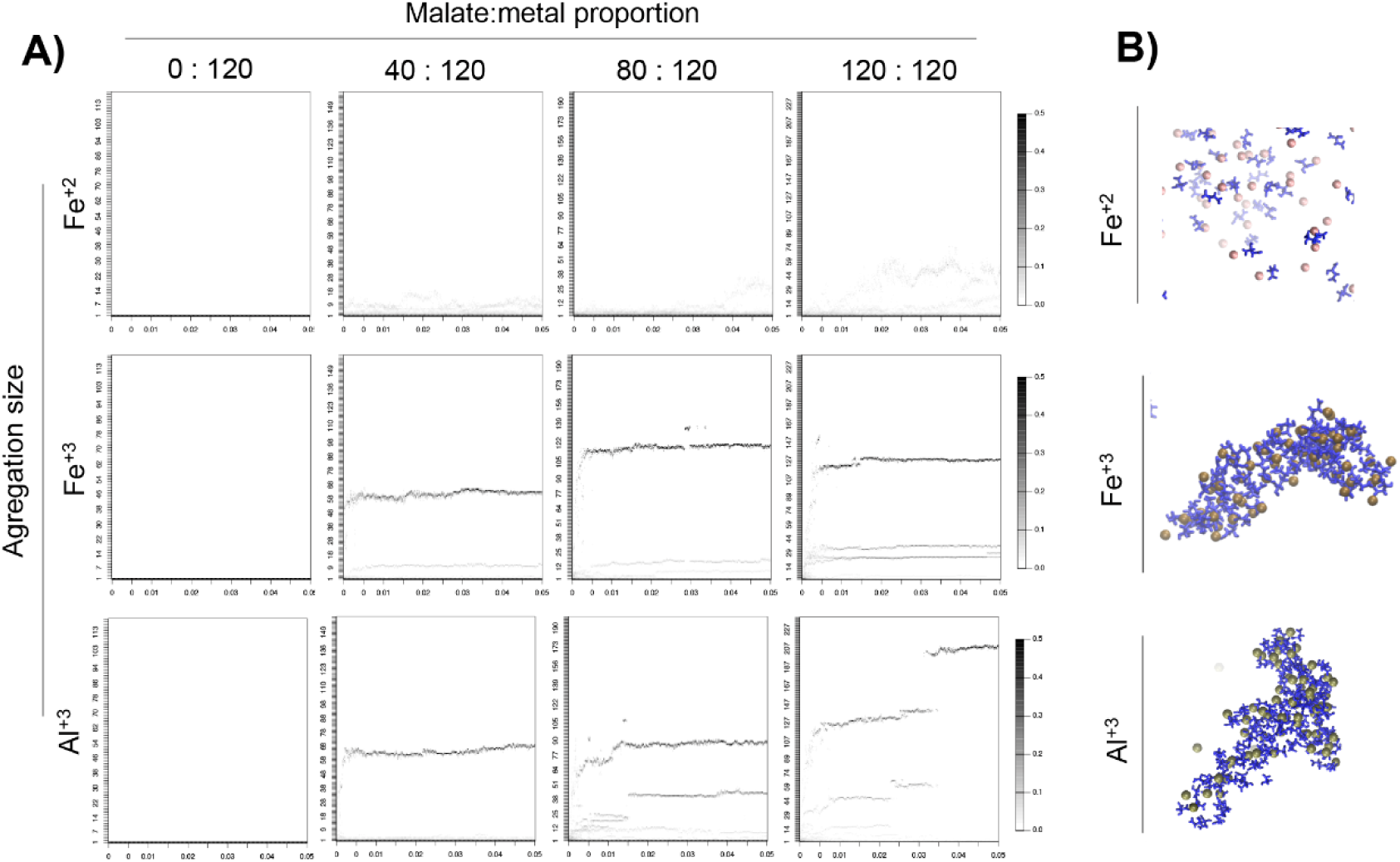
Molecular dynamic calculations revealed the malate-chelating effect that induces Fe^3+^ aggregation. 120 molecules of each metal (A-D) Fe^2+^, (E-H) Fe^3+^ and (I-L) was simulated (see Materials and Methods) with 0, 40, 80 and 100 molecules of malate, respectively. Structural representation at the end (50 microseconds) of simulation between the higher malate concentration (120 molecules) and (M) Fe^2+^, (N) Fe^3+^ and (O) Al^3+^, respectively.

**Supplementary Figure 6.**
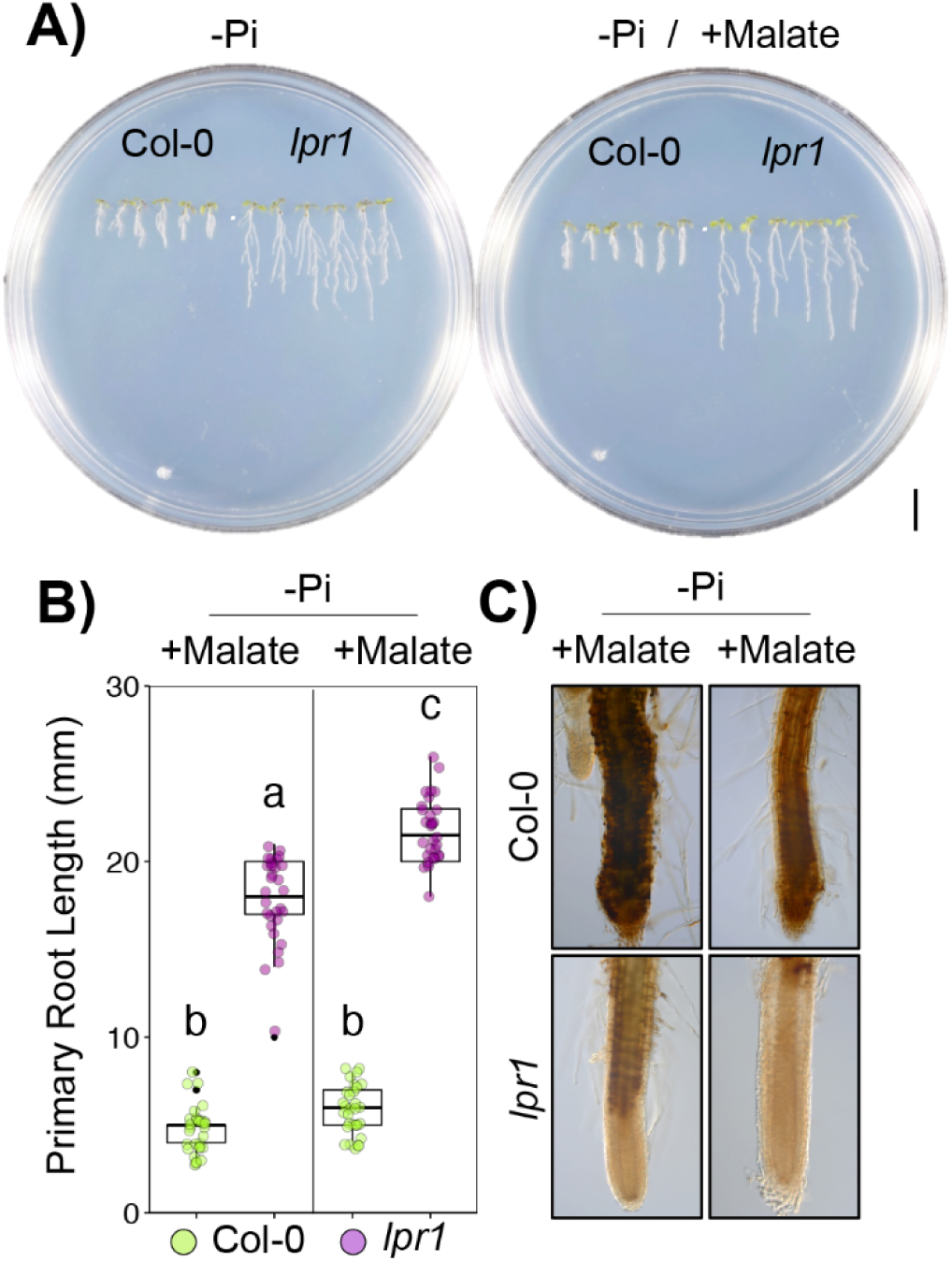
Malate supplementation of low Pi medium did not rescue the long root phenotype of *lpr1* mutants. (A) Phenotypes of Col-0 (WT) and *lpr1* seedlings grown under -Pi conditions (left) and -Pi medium supplemented with 1 mM malate (-Pi+M) (right) conditions 10 dag. (B) Primary root length of 10 dag WT and *lpr1* seedlings grown under -Pi conditions and -Pi+M conditions. (C) DAB-Perls iron staining of roots from Col-0 and *lpr1* seedlings grown under -Pi and -Pi+M conditions 10 dag.

**Supplementary Figure 7.**
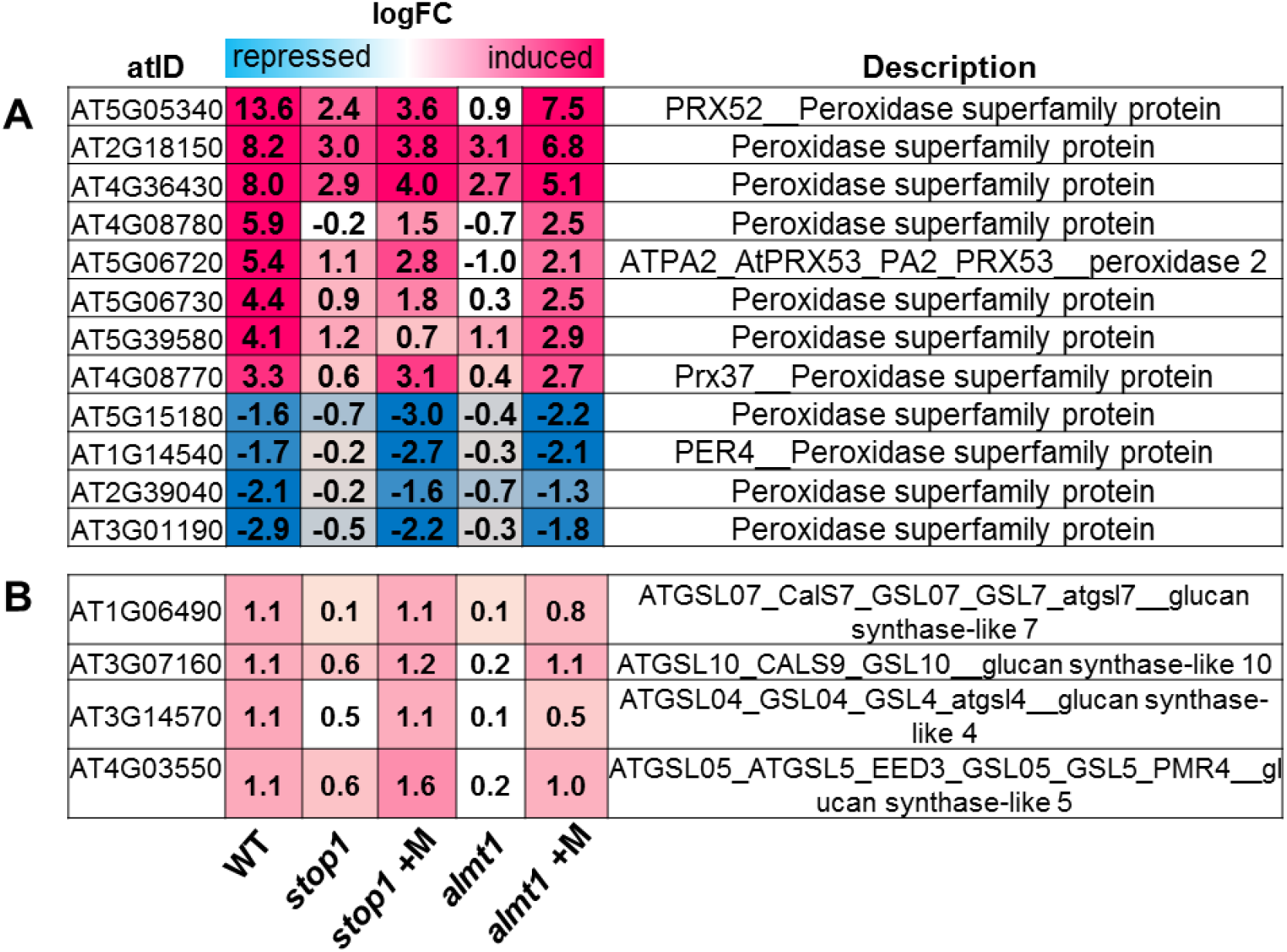
Malate treatment rescues the expression of peroxidase and callose synthase genes in Pi-deprived *stop1* and *almt1* seedlings. The locus ID (atID), description and the expression values (logFC) of (A) *PEROXIDASE* and (B) *CALLOSE SYNTHASE* genes are presented for Pi-deprived seedlings (WT, *stop1* and *almt1*) and malate-treated Pi-deprived seedlings (*stop1*+ M, *almt1*+M). Non colored values mean that the gene is not differentially expressed (FDR>.05) in the respective genotype/conditions.

**Supplementary Figure 8.**
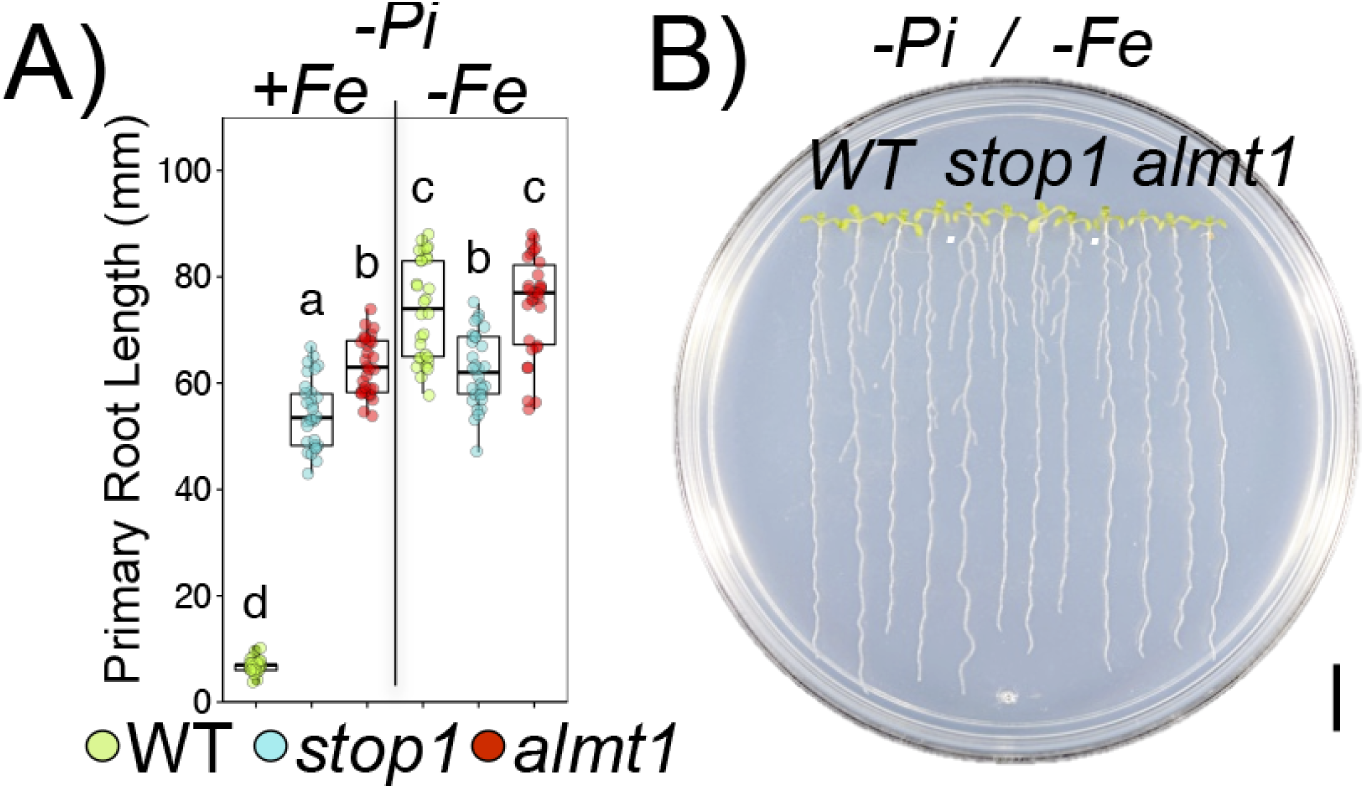
Iron-dependence of the inhibition of primary root under low Pi conditions. (A) Primary root length of 10 dag WT, *stop1* and *almt1* seedlings grown In - Pi media (-Pi+Fe) and –Pi media lacking Fe (-Pi-Fe; See Materials and Methods). Green, blue and red dots depict WT, *stop1* and *almt1* individuals (n=30 from 3 independent experiments), respectively. (B) Phenotypes of 10 dag WT, *almt1* and *stop1* seedlings grown under -Pi-Fe conditions. Scale bar equals 1 centimeter (cm).

